# Targeting stiffness-dependent YAP/TAZ restores angiogenesis dynamics impaired by ALK1 knockout *in silico*

**DOI:** 10.1101/2025.09.26.678712

**Authors:** Margot Passier, Sandra Loerakker, Tommaso Ristori

## Abstract

Hereditary Hemorrhagic Telangiectasia (HHT) is a currently uncurable genetic disorder caused by loss-of-function mutations in the ALK1-BMP9 pathway, leading to dysregulated angiogenesis and consequential vascular malformations. Recent experiments also implicate the mechanotransducers YAP/TAZ in HHT pathology. However, how YAP/TAZ stiffness sensitivity and signaling activity contribute to aberrant HHT angiogenesis remains poorly understood.

Here, we extended our previous computational framework of stiffness-mediated YAP/TAZ-VEGF-NOTCH crosstalk to account for ALK1 signalling and predict the resulting angiogenic temporal dynamics. Our simulations predicted that ALK1 knockout impairs NOTCH activation, slowing endothelial phenotypic selection and shuffling while enhancing filopodia activity, features corresponding with hypersprouting. These effects were most pronounced in low stiffness environments, consistent with the previously observed prevalence of HHT vascular malformations in low stiffness organs. Importantly, the temporal dynamics of endothelial phenotypic selection and shuffling, as well as key protein activity levels, were partially restored by direct or cytoskeleton-mediated inhibition of YAP/TAZ resulting from increased NOTCH activation.

These computational findings offer more mechanistic insight into the signalling pathways and temporal dynamics of endothelial phenotypic selection underlying HHT vascular anomalies, and suggest that targeting YAP/TAZ and endothelial stiffness sensitivity may offer a promising therapeutic strategy to restore physiological angiogenesis.

**Author Summary:** Mechanical cues such as extracellular matrix stiffness, sensed by endothelial cells lining blood vessels, play a critical role in angiogenesis, the process of blood vessel formation from pre-existing vessels. Understanding the impact of these cues on angiogenesis in health and in diseased conditions could help guide new treatments for angiogenesis-related disorders, such as Hereditary Hemorrhagic Telangiectasia (HHT). In HHT, genetic mutations disrupt normal vessel development, leading to malformations that can rupture and have detrimental consequences. Here, we developed a computational model to investigate the effects of HHT-associated genetic mutations on angiogenic signaling of endothelial cells exposed to different stiffnesses. Simulations predicted that the mutation impairs endothelial phenotypic selection and shuffling, necessary for physiological angiogenesis. The mutation effects to be largest in soft organs and identified the mechanotransducers YAP/TAZ as possible targets to restore physiological signaling. Therefore, with this model, we have created a platform to simulate endothelial cell behavior in HHT patients, allowing us to explore mechanosensitive pathways as potential targets potentially extending future treatment options.

## Introduction

Hereditary hemorrhagic telangiectasia (HHT) is a currently uncurable genetic disease, affecting approximately 1 in 5000 people, characterized by the development of vascular malformations(1–3). The most commonly observed malformations are telangiectasia, small widened vessels in the mucosa and superficial skin layers(2,4). These fragile vessels are prone to rupture, often resulting in recurrent bleeding and anemia(5). Moreover, patients frequently develop arteriovenous malformations (AVMs), direct connections (shunts) between arteries and veins replacing the physiological capillary bed(2,6,7). Such AVMs predominantly form in the lungs, liver, and brain(2,4,8), potentially resulting in serious complications such as rupture and consequential bleeding or organ failure. In most cases (over 92%), HHT is caused by mutations affecting the activin-receptor like kinase 1 (ALK1) – Bone Morphogenic Protein 9 (BMP9) signaling pathway(2,9–14). Increasing evidence shows that these mutations dysregulate angiogenesis, the process of new blood vessel formation from pre-existing vessels(15), thereby leading to the vascular malformations mentioned above. However, the underlying mechanisms are still unclear, hindering the development of definitive cures.

Under physiological conditions, ALK1-BMP9 signaling regulates Vascular Endothelial Growth Factor (VEGF)- NOTCH signaling crosstalk, a key regulator of sprouting angiogenesis(16). Briefly, VEGF kickstarts sprouting angiogenesis by activation of the VEGF-Receptor 2 (VEGFR2), which induces previously quiescent endothelial cells (ECs) to adopt the tip and stalk phenotypes. Specifically, VEGFR2 activation upregulates filopodia formation and expression of the Notch ligand Delta-like ligand 4 (DLL4), thereby inducing a migratory tip phenotype. Simultaneously, DLL4 binds to and activates the NOTCH1 receptor on neighboring cells, causing downregulation of VEGFR2 and upregulation of VEGF-Receptor 1 (VEGFR1, a decoy receptor for VEGF(17–19)). The VEGF-sensing ability is therefore downregulated in these NOTCH1-activated cells and they are induced to acquire a stalk cell phenotype(20,21). This process is highly dynamic, as cells continue to rearrange their position and to switch phenotype(22,23). Knockout (KO) of the BMP9-receptor ALK1 (ALK1 KO), the causal mutation of the HHT genetic subtype HHT2, disrupts this balance of angiogenic stimuli(24). It leads to decreased activity of NOTCH target genes HES and HEY, and consequentially downregulated VEGFR1 activity and upregulated VEGFR2 activity(25). This causes overactivity of pro-angiogenic stimuli through active VEGFR2, which is thought to be one of the underlying mechanisms causing the vascular anomalies observed in HHT2 patients.

In addition to directly affecting NOTCH targets, ALK1 activation has recently been shown to upregulate the expression of Lunatic Fringe (LFNG), a glycosyltransferase that enhances NOTCH1-DLL4 binding affinity and activation(26–28). Al Tabosh et al. identified LFNG as one of the few genes consistently dysregulated in newborn HHT patients carrying the ALK1 heterozygous mutation(26). This suggests a central role of LFNG in HHT and its corresponding aberrant angiogenesis. In support of this, LFNG inhibition has been shown to cause hypersprouting(27). Furthermore, Ristori et al. have recently shown that BMP9 regulates NOTCH1 and sprouting angiogenesis by increasing the expression of LFNG(29). Hence, these studies implicate LFNG as one of the key players in HHT2.

Another recent study has shown that mechanotransducers YAP/TAZ also play a major role in HHT2(2,6). YAP/TAZ, transcriptional co-activators of the Hippo pathway, are mechanosensors that are mainly located in the cytoplasm for cells on substrates with relatively low stiffness, while their nuclear localization (and thus regulation of gene expression) increases with increasing substrate stiffness(30–33). Nuclear translocation of YAP/TAZ is initiated by integrin binding, followed by multiple cytoskeletal processes culminating in stress fiber formation and consequential nuclear deformations allowing YAP/TAZ nuclear translocation(34–36). VEGF also influences this process; VEGFR2 activation upregulates focal adhesion kinase (FAK) phosphorylation, involved in the stress fiber formation and nuclearization of YAP/TAZ (37,38). Interestingly, inhibition of YAP/TAZ has recently been shown to prevent AVM formation in ALK1 KO mice via rescued EC polarization sensitive to blood flow(6), indicating that YAP/TAZ are involved in HHT AVMs. However, how YAP/TAZ sensitivity to stiffness and its crosstalk with angiogenic signaling pathways fit into this picture are currently unclear.

Previous experiments have shown that YAP/TAZ downregulate the expression of LFNG and DLL4(39,40). By using computational simulations, we have recently shown that this YAP/TAZ-NOTCH1 crosstalk could explain the angiogenic response to stiffness(41). In particular, we found that, through these interactions, YAP/TAZ is a temporal regulator of EC phenotypic selection speed(41) and thus vascular topology(42,43). In the present study, we hypothesized that this crosstalk might be essential in HHT2 as well. More specifically, we computationally investigated whether the YAP/TAZ-sensitivity and interaction with the NOTCH pathway can explain and potentially be targeted to rescue the abnormal EC behavior and phenotypic selection observed in response to ALK1 KO(29,44), specifically in tissues prone to AVM formation in HHT2, such as the liver, characterized by relatively low stiffness(1,3,24,45–50).

In the field of angiogenesis, there have been many advances resulting from computational models addressing key aspects of EC behavior(23,29,42,43,51–64). An integration of simulations and experiments for example, led to the discovery of EC phenotypic and positional shuffling(22,23), and to the observation that the EC phenotypic selection and shuffling rates influence vascular network topology. For instance, faster patterning dynamics are associated with the formation of denser networks(42). For mechanisms also involved in HHT specifically, computational approaches have been adopted to establish the importance of LFNG in BMP9-regulated vascular patterning(29), and to explore VEGFR1-VEGFR2 competition, elucidating its impact on sprout directionality(65). Moreover, computational studies focused on HHT2 mainly addressing the characteristics of mutations and diagnostics, contributed to the emerging view that HHT2 might be substantially underdiagnosed(66).

In this study, we developed a computational model to gain more insight into the role of YAP/TAZ stiffness sensitivity and its effect in HHT2. Specifically, we extended our previous YAP/TAZ-VEGF/NOTCH1 framework for stiffness-regulated tip/stalk pattern formation based on ordinary differential equations (ODEs)(35,41,58) by: (i) including the effects of BMP9-ALK1 signaling on the VEGF-NOTCH1 crosstalk(29); (ii) differentiating between VEGFR1 and VEGFR2(17,19,22,67); and (iii) accounting for VEGF-induced FAK activation(37,38,68,69). The ensuing simulations, focusing on EC phenotypic selection, allowed us to dive into the effects of YAP/TAZ stiffness sensitivity on ECs that are affected by the ALK1 KO mutation. Our simulations indicated that ALK1 KO significantly altered signaling dynamics, leading to decreased NOTCH activity (especially on lower stiffnesses), causing decreased lateral inhibition, thereby slowing down phenotypic selection. Furthermore, we found that ALK1 KO ECs have an impaired ability to switch phenotypes and studied knock down of cytoskeletal elements as possible new targets in HHT2. We found that YAP/TAZ inhibition could partially rescue physiological behavior, such as phenotypic selection time, but not completely, potentially because LFNG expression was not completely restored. We highlight the potential of the identified signaling network and developed computational model as a screening and simulation tool for conducting more *in silico* experiments.

## Results

### The extended model captures VEGF- and stiffness-dependent YAP/TAZ nuclearization and tip/stalk temporal dynamics

We first verified whether our computational model, extended with the VEGF-FAK interaction and VEGFR1-2 distinction, could reproduce the experimentally-observed YAP/TAZ nuclearization dependent on VEGF activity and stiffness(37,70), as well as the trends in angiogenic patterning behavior in response to stiffness observed in our previous computational study(41). Consistent with our previous simulations(41), the extended model predicted that YAP/TAZ nuclearization increased in response to stiffness following a logistic shape (Fig. 1A), resulting from the stiffness-mediated increase in phosphorylated FAK (equation 1. methods). Due to the model modifications, the extended model could successfully capture the increased YAP/TAZ nuclearization caused by increased VEGF exposure too, as a consequence of VEGF-mediated upregulation of phosphorylated FAK (Fig. 1A, equation 1, methods). The simulations also showed the biphasic relationship between patterning speed and stiffness, predicting the highest patterning rate(71,72) for an intermediate stiffness (Fig. 1B,G), as well as increased filopodia activity for stiffer substrates (Fig. 1C,G), in agreement with our previous simulations(41). These trends are the result of increased nuclear YAP/TAZ content upon increased stiffness (Fig. 1A), leading to more DLL4 and LFNG inhibition (equations S8, S12-13, Fig. 1E-F). DLL4 expression was however predicted to recover for stiffnesses above 20 kPa (Fig. 1E), as also observed in our previous study(41). This resulted from the YAP/TAZ-mediated decrease in LFNG, causing impaired NOTCH1 activation and therefore VEGFR2 upregulation, causing all cells to become tip cells (Fig. 1D) and express more DLL4 (Fig. 1E). These results show that the extended model can predict not only the effects of stiffness on YAP/TAZ and NOTCH1-mediated tip/stalk patterning, but also the effects of VEGF on YAP/TAZ.

**Figure 1:**
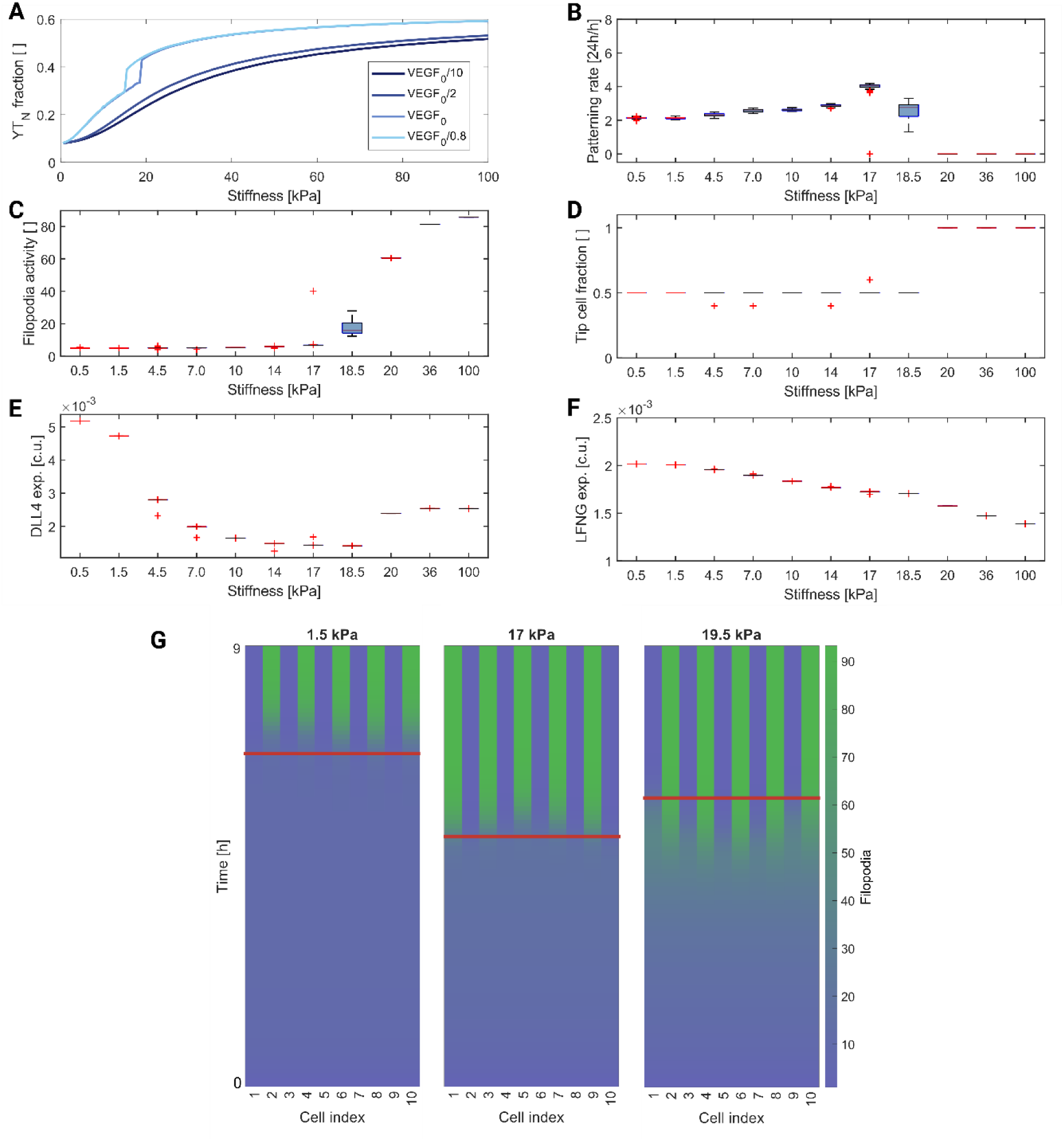
The extended model captures the biphasic response to stiffness and important trends of the original model. (A) Mean trends of YAP/TAZ nuclear fractions for different amounts of VEGF. For the VEGF concentration used to instigate patterning VEGF_0_, the patterning rate (24h/patterning time, B), filopodia activity prior to pattern formation (C), fraction of tip cells (D), expression of DLL4 (E) and expression of LFNG (F) are shown. All simulations were repeated 25 times, with the boxes in B-F depicting quartiles 2 and 3, the red lines indicating the medians, the whiskers indicating the min- and maxima and the red plusses indicating the outliers. For (A), only the mean is plotted as the standard deviation was so small (and invisible) that we omitted it. (G) Representative images of filopodia activity over time, for three different stiffnesses (1.5, 17 and 19.5 kPa). The red lines indicate the timepoint at which the pattern is considered established.

### YAP/TAZ inhibition can partially rescue phenotypic selection behavior in ALK1 KO ECs

Next, we investigated the effects of ALK1 KO and YAP/TAZ KO on the temporal dynamics of tip/stalk patterning. Liver stiffness (approximately 5 kPa(46,47,73)) was taken as a reference, given the frequent occurrence of hepatic AVMs in HHT2(1,3,24). Consistent with Park et al.(6), we considered three groups: wild-type ECs, ECs with ALK1 KO (modelled through their effect on HES/HEY and LFNG, Fig. 7), and ECs with both ALK1 KO as well as YAP/TAZ KO.

Phenotypic selection slowed down upon ALK1 KO (Fig. 2A, D). In addition to the ECs needing almost twice as much time to establish the tip/stalk pattern (Fig. 2A,D), filopodia activity (Fig. 2A,E) as well as DLL4 expression prior to patterning (Fig. 2F) increased significantly, suggesting tip-like behavior, indicative of EC hyperactivity and hypersprouting (Fig. 2A)(42,43). This agrees with experimentally observed behavior of ALK1 KO ECs(44,74). In the simulations, this resulted from the marked increase in VEGFR2 activity (Fig. 2G), consistent with experimental findings(25). Interestingly, it was not only the activity of tip cell indicators (filopodia and DLL4) that increased, but also NOTCH1 activity (or NICD), indicative of stalk cells, prior to patterning (Fig. 2C,H). This indicates a reduced cellular ability to perform lateral inhibition. Although average NOTCH1 activity prior to patterning increased upon ALK1 KO (Fig. 2C,H), NOTCH1 activity in patterned stalk cells appeared to be lower compared to that of wild-type ECs (Fig. 2C), possibly due to decreased levels of LFNG leading to decreased DLL4-NOTCH1 binding affinity (Fig. 2I). YAP/TAZ nuclear fraction was also seen to increase upon ALK1 KO, in agreement with experimental findings(6), especially prior to pattern formation (Fig. 2B,J). VEGFR1 expression was not much affected by ALK1 KO (Fig. 2K). Overall, the ALK1-KO simulations suggested that EC hypersprouting experimentally observed in ALK1-KO angiogenesis(44) could result from delayed tip-cell inhibition and thus delayed tip-stalk pattern formation accompanied by high EC filopodia activity.

**Figure 2.**
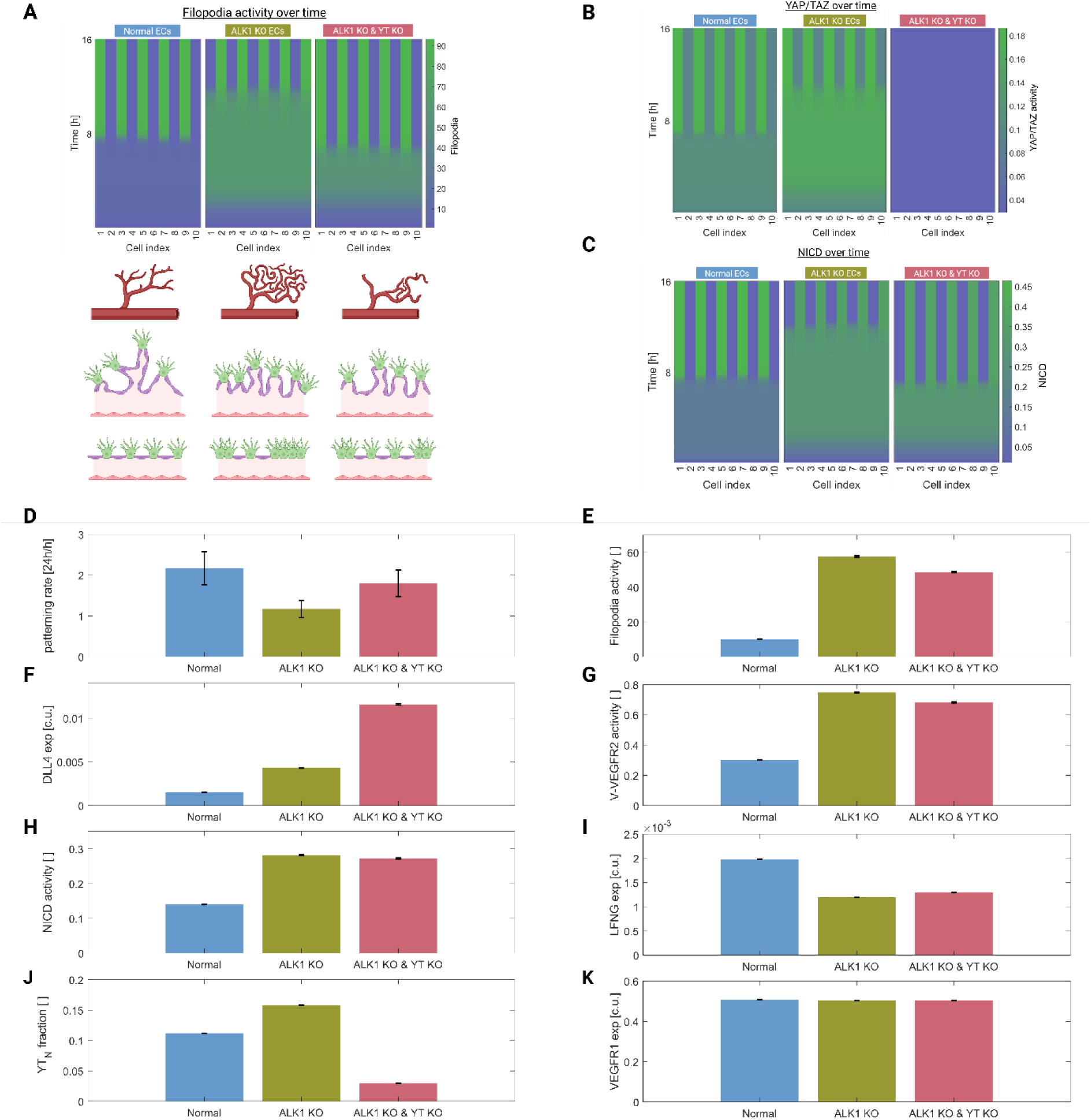
Simulations of ALK1 KO ECs indicate a decreased patterning rate and increased filopodia activity, indicative of hypersprouting. Panels A-C show time courses of filopodia activity (A), YAP/TAZ nuclearization (B) and NOTCH activity (NICD) (C) during the first 16 hours after being provided with a VEGF stimulus to instigate patterning. (A) provides an additional schematic of the predicted vascular topology, indicative of optimal patterning with physiological sprouting (left), hypersprouting (middle), and an in-between case (right). ALK1 KO ECs exposed to liver stiffness (yellow bars) show a decreased patterning rate (D), LFNG expression (I), and minimal decrease in VEGFR1 expression (K), while filopodia activity (E), DLL4 expression (F), VEGFR2 activity (G), NICD (H) and YAP/TAZ nuclearization (J) all increased. Except for DLL4 (F), all behaviors partially returned to wild-type signaling levels (light blue bars) upon YAP/TAZ inhibition (pink bars). All simulations were repeated 25 times, showing the mean with standard deviation represented by the error bars (D-K). For E-K, mean activity and expression were determined by averaging the amounts over the timepoints prior to pattern formation. The schematic in (A) was created using Biorender.com

In response to ALK1 KO, the expression of NOTCH1 target genes HES/HEY was also seen to increase prior to pattern formation (Fig. 3A), in apparent disagreement with prior experiments(29) and the observed increase in VEGFR2 (Fig. 2G). However, analysis of HES/HEY activity over time revealed that, the observed increase in HES/HEY expression for ALK1 KO ECs prior to patterning, appeared only after approximately 1.5 hours. During the first 1.5 hours, HES/HEY expression in ALK1 KO ECs was lower compared to wild type (Fig. 3A). This time dynamics can be explained by the fact that, the initial decrease in HE expression led to lower VEGFR2 inhibition and VEGFR1 upregulation, causing VEGFR2 activity to slowly increase over time. The increased VEGFR2 activity led to upregulation of filopodia formation, more VEGF sensation (Fig. 3B) and as a consequence, even more VEGFR2 activity (establishing a positive feedback loop). Increased VEGFR2 activity also upregulated DLL4 expression, leading to increased DLL4-NOTCH1 binding and, finally, increased NICD levels and thus HES/HEY expression levels too (Fig. 3).

**Figure 3.**
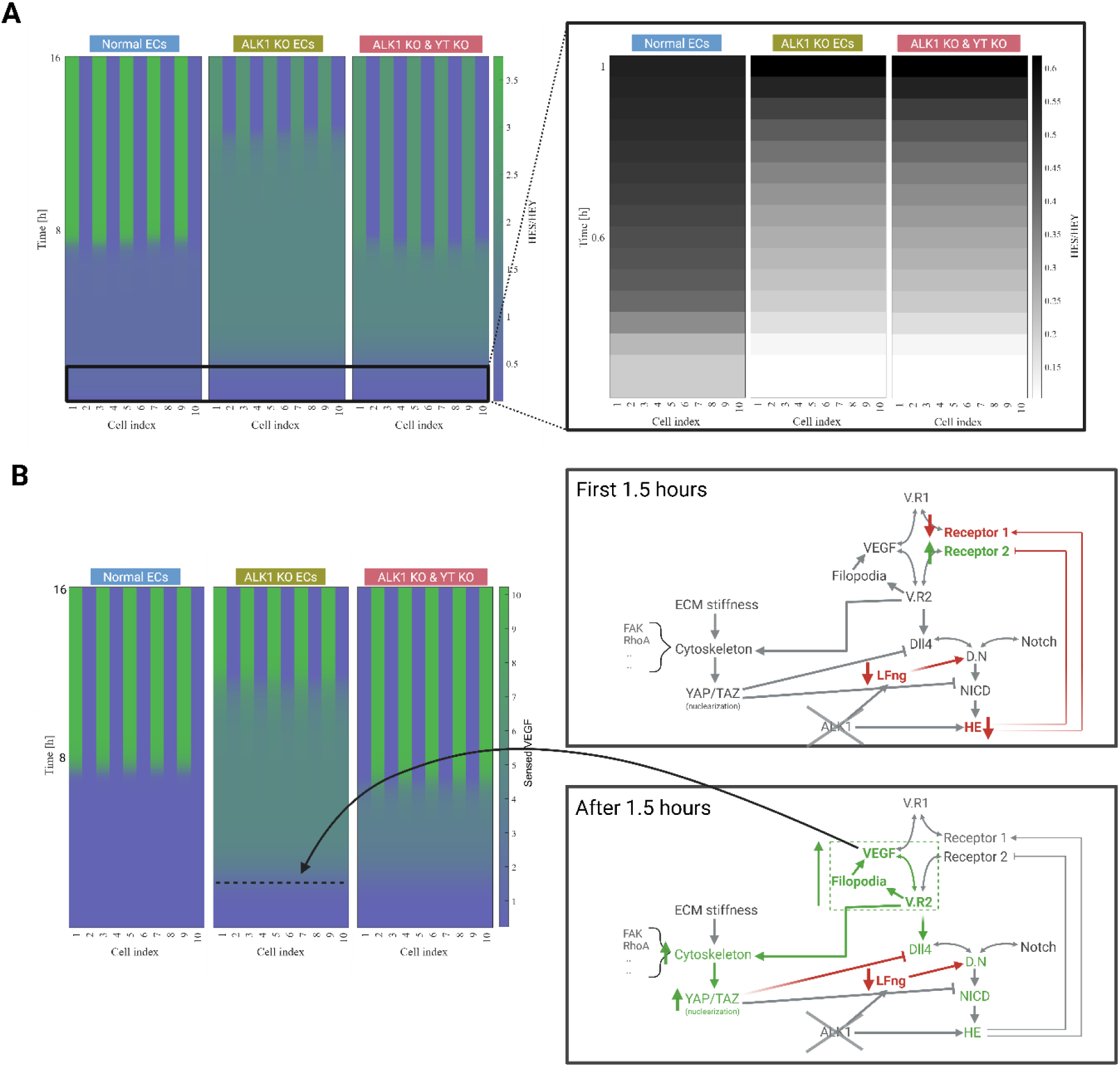
HES/HEY expression initially decreases upon ALK1 KO, but eventually increases due to over-active VEGFR2 signaling. (A) Representative time course of HES/HEY levels for wild-type ECs (left panels, light blue heading), ALK1 KO ECs (middle panels, yellow heading) and ALK1 KO + YAP/TAZ KO ECs (right panels, pink heading) for 16 hours. The right window of (A) depicts a zoom-in of the temporal dynamics of HES/HEY expression over the first hour. (B) Time course analysis of VEGF levels, which start to increase after HES/HEY levels of ALK1 KO ECs have increased, resulting from the positive feedback loop consisting of VEGFR2-filopodia-VEGF, depicted in the schematic in the right window. The schematic in (B) was created using Biorender.com

Park et al. experimentally demonstrated that inhibition of YAP/TAZ can prevent AVM formation(6). Consistent with their experiments, we predicted an increase in YAP/TAZ nuclearization upon ALK1 KO (Fig. 2J). YAP/TAZ inhibition led to a major increase in patterning rate compared to the ALK1 KO case without YAP/TAZ inhibition, bringing it in much closer alignment with wild-type patterning times (Fig. 2A,D). Upon YAP/TAZ KO, LFNG expression in ALK1 KO ECs increased slightly and DLL4 expression increased two-three fold, therefore leading to stronger lateral inhibition potential (Fig. 2F,I). Moreover, the marked increase in VEGF levels in ALK1 KO ECs which resulted from the positive VEGFR2-filopodia-VEGF feedback loop, was significantly lower upon YAP/TAZ KO (Fig. 3B). Consequently, overall filopodia and VEGFR2 activity prior to pattern formation decreased and cells established the tip/stalk pattern quicker (Fig. 2E,G). Therefore, despite YAP/TAZ KO not being able to restore all protein levels in the phenotypic selection process, it could nevertheless rescue the tip/stalk patterning rate and thus limit hypersprouting due to increased LFNG and DLL4 expression (Fig. 2A). These simulations therefore indicate that YAP/TAZ inhibition may lead to more physiological angiogenesis by mediating the VEGF-NOTCH crosstalk via LFNG and DLL4 regulation.

### YAP/TAZ KO can partially rescue EC phenotypic shuffling impaired by Alk1 KO

In addition to the tip/stalk pattern selection at the onset of sprouting, the final vascular network is strongly influenced by the tip/stalk shuffling rate(22,23,75). Our recent *in vitro* experiments have shown that LFNG KD cells are more prone to shuffle positions within the sprout(29). Given the interaction of ALK1 and YAP/TAZ for the regulation of LFNG, following a similar approach as in Venkatraman et al.(58), we here investigated the ability of ALK1 KO cells to switch phenotypically (i.e. going from tip to stalk or vice versa) upon a positional change, and checked whether YAP/TAZ KO could partially counteract the ALK1-KO effects. To mimic a VEGF gradient, we initially exposed two ECs to 100% and 90% of the reference VEGF cue, ensuring the establishment of a distinct cell fate for both cells. After 24 hours, to mimic a positional switch, the external stimuli (DLL4 and VEGF) were inverted (Fig. 4A). Given the possible variations in and the effects of the steepness of the VEGF gradient for the resulting potential phenotypic switch, we varied the lower level of VEGF sensed (between 10 to 90% of the amount of VEGF the other cell senses, see methods, equation 11).

**Figure 4:**
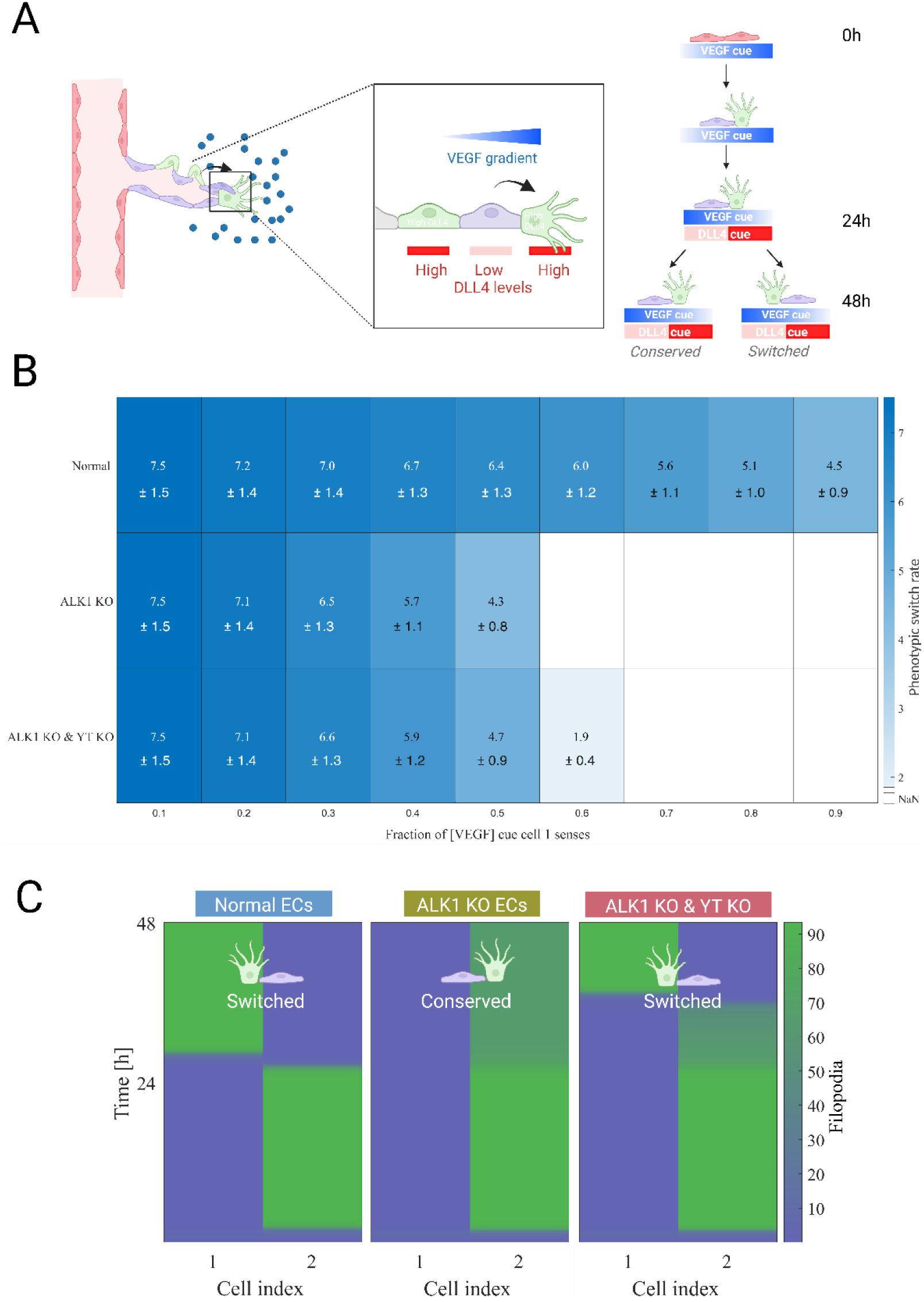
ALK1 KO ECs have impaired phenotypic shuffling ability that can be partially rescued by YAP/TAZ inhibition. (A) Schematic of the different simulated phases mimicking positional switching during angiogenic sprouting. A tip and stalk cell switch position, thereby switching the environmental cues that they sense. This is mimicked by first (0-24h) exposing the cells to different VEGF levels, ensuring the right cell becomes a tip cell. After 24h, the environmental cues are switched, mimicking a positional change, leading to either conserved or switched phenotypes. (B) Average shuffling rate with standard deviation (simulations conducted 25 times), for the range of different VEGF ratios that was simulated. White squares are indicative of conserved phenotypes (i.e. no switch, which are largely present in the ALK1 KO row). (C) Time course analyses of filopodia activity for cell 2 sensing 60% of the amount of VEGF sensed by cell 1, showing switched phenotype for wild-type ECs (left panel), conserved phenotypes for ALK1 KO ECs (middle panel) and delayed switched phenotypes for ALK1 KO & YAP/TAZ KO ECs (right panel). The schematics in (A) and (C) were created using Biorender.com

The model predicted that, following a positional switch, wild-type ECs were able to relatively quickly switch phenotype, independent of the difference in the amount of sensed VEGF (Fig. 4B,C). In contrast, ALK1 KO cells switched phenotype at a generally lower rate compared to wild type, and only if exposed to VEGF values differing at least a factor 2, representing relatively steep VEGF gradients. This is consistent with the impaired ability of ALK1 KO cells to repress the tip phenotype in their neighbors due to decreased LFNG expression. This behavior was partially rescued by YAP/TAZ KO. Cells switched faster for cases in which one of the cells sensed 30-50% of the amount of VEGF the other cell was sensing. More importantly, when one cell sensed 60% of the amount of VEGF sensed by the other cell, the phenotypic switch that was inhibited by ALK1 KO was instead rescued by YAP/TAZ KO. Representative simulations showing the latter case are provided in Fig. 4C, depicting filopodia activity over time. Even though the ALK1 KO ECs showed a clear decrease in the amount of filopodia for the initial tip cell upon the shuffling cue, the initial stalk cell was unable to repress the tip phenotype in its neighbor due to low LFNG and thus weakened lateral inhibition, causing the cells to maintain their original phenotypes (Fig. 4C). YAP/TAZ KO could counteract this effect; in these simulations, the positional shuffling was followed by an initial decrease in filopodia expression in the original tip cell, prior to this cell adopting the stalk phenotype and the other cell taking on a tip phenotype (Fig. 4C). This was probably caused by increased LFNG, partially restoring lateral inhibition potential. These simulations therefore indicate that ALK1 KO ECs maintain a high migratory behavior (indicated by high filopodia activity) also when they lose the tip position, consistent with increased positional switching observed in LFNG KO experiments(29). Moreover, the simulations suggest that YAP/TAZ KO may partially rescue the physiological phenotypic switching potential.

### ALK1 KO primarily affects EC behavior in low stiffness regimes

YAP/TAZ nuclear translocation and NOTCH activity are known to respectively increase and decrease in response to stiffness(30,40,70,76). Therefore, motivated by the interaction between YAP/TAZ and NOTCH as well as NOTCH with ALK1, we next examined the dependence of ALK1 KO EC behavior on different stiffnesses. For low stiffnesses, the model predicted decreased patterning rates for ALK1 KO compared to wild type. Patterning of ALK1 KO ECs ceased for stiffnesses higher than 6 kPa, a threshold much lower compared to wild-type ECs (19 kPa, Fig. 5A). For stiffnesses above 19 kPa, wild type and ALK1 KO EC behavior was relatively similar, although wild-type ECs were a bit slower to all adopt the tip phenotype (Fig. 5I,J). Additionally, the model predicted increased filopodia activity prior to pattern formation, especially for stiffnesses lower than 19 kPa (Fig. 5B). The combination of these two behaviors in ALK1 KO ECs for low stiffnesses (until ceased patterning at 6 kPa) corresponds to hypersprouting, consistent with the behavior observed for ALK1 KO ECs on liver stiffness (Fig. 2).

**Figure 5.**
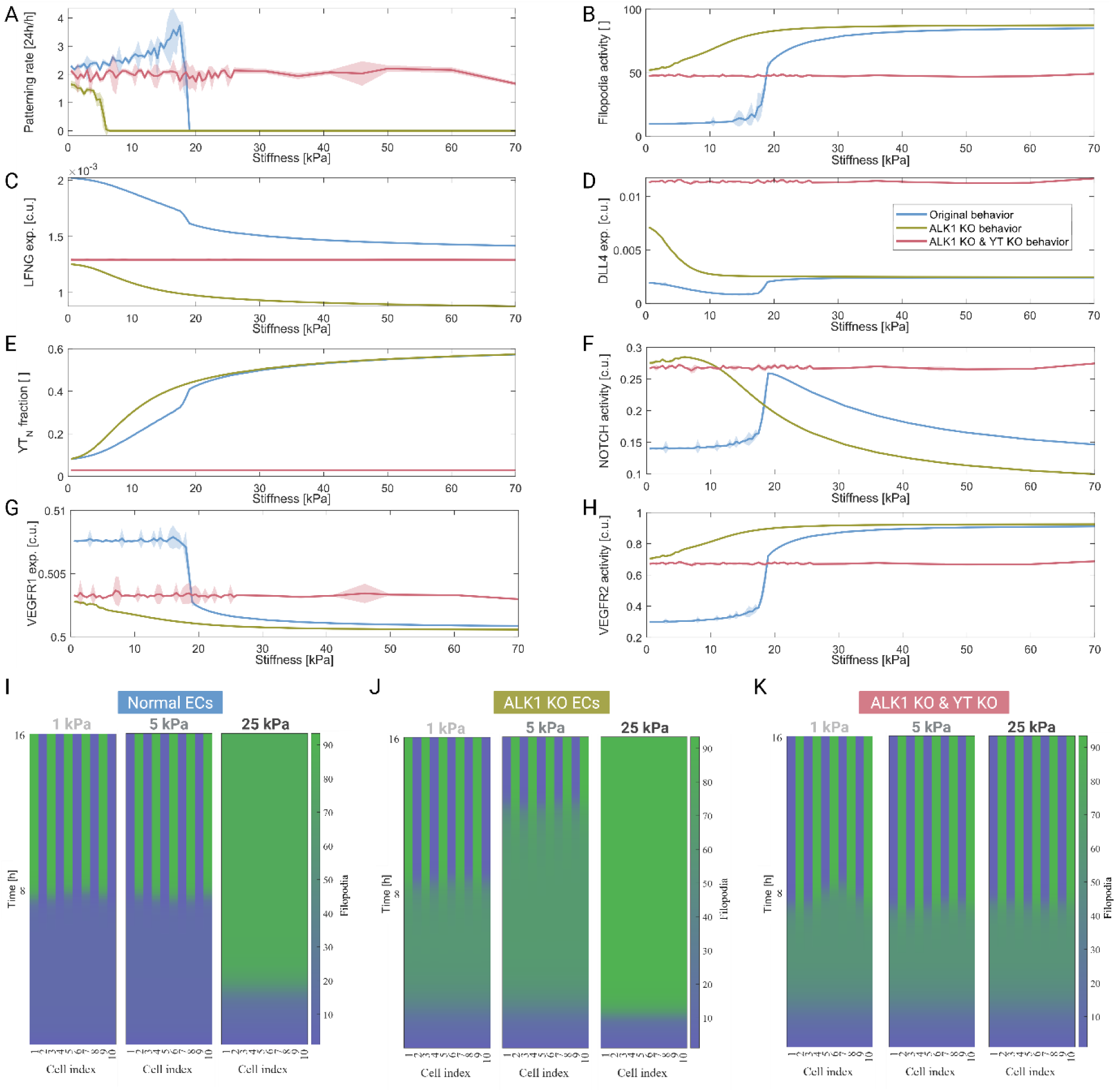
YAP/TAZ inhibition can rescue EC behavior in ALK1 KO cases, which is mainly affected in low stiffness regimes. The simulations predicted increased patterning rate (A), filopodia activity (B), DLL4 expression (D), YAP/TAZ nuclear fraction (E), NOTCH activity (F) and VEGFR2 activity (H) for ALK1 KO ECs (yellow) compared to wild-type cells (light blue), while LFNG expression (C) and VEGFR1 expression (G) decreased. Protein levels remained stable for all stiffnesses upon ALK1 KO & YT KO (pink graphs). All simulations were repeated 25 times, here showing the average with standard deviation in shaded color. For panels B-H, mean activity or expression levels were determined by averaging over the timepoints before patterning occurred (see methods). Representative time course analyses are shown for filopodia activity at three different stiffnesses; 1 kPa, 5 kPa and 25 kPa, for normal ECs (I), ALK1 KO ECs (J) and ALK1 KO with YAP/TAZ KO ECs (K). Mainly normal and ALK1 KO ECs show increasing filopodia activity with increasing stiffness.

Part of this slower patterning response can be explained through LFNG, which showed significantly lower expression in ALK KO ECs throughout the whole range of simulated stiffnesses, thereby weakening lateral inhibition (Fig. 5C). For both wild-type and ALK1 KO ECs, LFNG (and DLL4, Fig. 5D) expression decreased for increasing stiffness, resulting from increased nuclear fractions of YAP/TAZ (Fig. 5E). Similar to the behavior described for filopodia activity, DLL4 expression, the YAP/TAZ nuclear fraction, NICD and VEGFR2 activity all increased upon ALK1 KO for relatively low stiffness, most significantly below 19 kPa (stiffness for which wild-type ECs ceased patterning) (Fig. 5D-F,J). The concurrent elevation of these factors can be explained by the positive feedback loop leading to amplified VEGFR2 activity and consequentially increased HE activity, as mentioned earlier (Fig. 3B). All behaviors, except for NOTCH activity, were predicted to get closer to wild-type levels for stiffnesses above 19 kPa, while NOTCH activity went below wild-type expression levels (Fig. 5F). This latter trend could result from weakened lateral inhibition due to a sharp decrease in LFNG, which we observed perhaps because of the steep increase in nuclear YAP/TAZ at approximately 19 kPa. The stiffness response of YAP/TAZ is believed to be extra sensitive in ALK1 KO ECs, due to its implementation as a Hill function (see equation S25) which led to a more rapid increase in these parameters under ALK1 KO conditions, as modulated by the VEGF-FAK interaction. Interestingly, the largest behavioral differences were all observed for the low stiffness regions, corresponding to stiffnesses associated with organs frequently affected by AVM formation in HHT2 patients(2,4,8,46,77,78).

Finally, we checked how YAP/TAZ KO affected behavior of ALK1 KO ECs for different stiffnesses. Not surprisingly, YAP/TAZ KO caused ECs to lose (most of) their stiffness-dependent behavior. As a result, this caused the expression of DLL4 and LFNG (Fig. 5C-E), and therefore VEGFR2, VEGFR1 and NOTCH activities (Fig. 5F-H), to remain at approximately the same level for all stiffnesses. This resulted in the patterning rate and filopodia activity pre-patterning to remain at stable levels over the whole range of stimulated stiffnesses (Fig. 5A,B,K), with levels similar to wild-type ECs exposed to low stiffness. This indicates that YAP/TAZ KO can (partially) rescue the physiological filopodia formation and tip/stalk patterning rate for ALK1 KO ECs, independently of stiffness.

### Knockdown of cytoskeletal elements decreases YAP/TAZ signaling and rescues patterning in ALK1 KO ECs for higher stiffnesses

Simulations with our original model suggested that cytoskeletal perturbations can regulate tip/stalk dynamics by modulating YAP/TAZ activity(41). Therefore, we next investigated whether perturbing cytoskeletal elements could rescue phenotypic selection (temporal) dynamics of ALK1 KO ECs. For all inhibitors except Cofilin, due to its inhibitory relationship with YAP/TAZ, YAP/TAZ KO rescued patterning in ALK1 KO ECs for stiffnesses above 6 kPa (Fig. 6A). YAP/TAZ KO even enabled patterning of ALK1 KO ECs for stiffnesses beyond which patterning normally ceased for wild-type ECs (Fig. 6A, 5A). The simulations indicated that targeting elements more upstream in the signaling cascade provided the largest effect (more closely mimicking that of direct YAP/TAZ targeting). Interestingly, no patterning was observed at 100 kPa for the simulation involving the myosin inhibitor, suggesting that myosin is a less potent target in the prevention of AVM formation. For most inhibitors, filopodia activity stabilized at relatively high levels, similar to that of ALK1 KO ECs for low stiffnesses (Fig. 6B).

**Figure 6.**
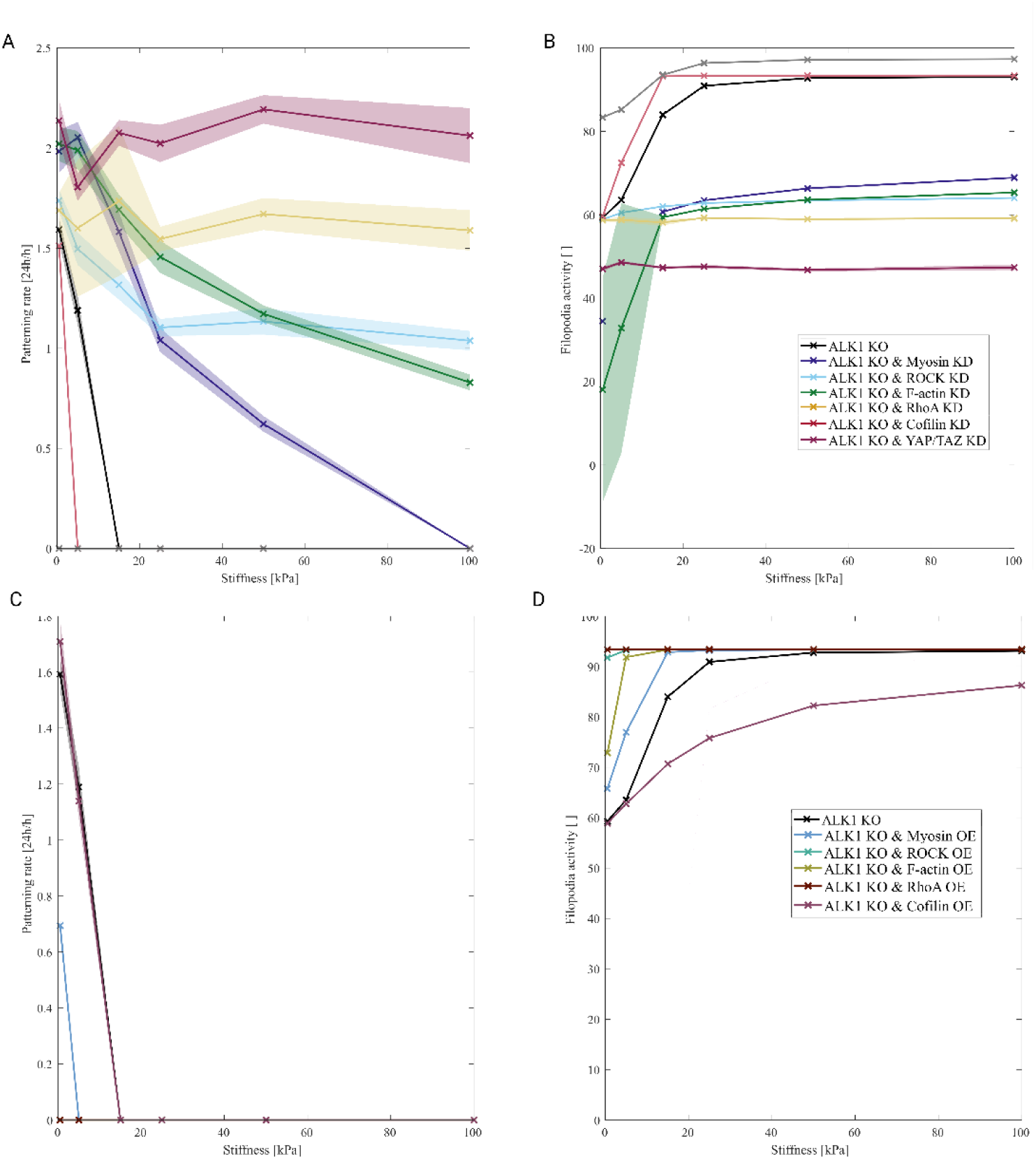
Knock down of cytoskeletal elements can rescue patterning and reduce filopodia activity in ALK1 KO ECs. Patterning rates (A) and average filopodia activity prior to pattern formation (B) for knock down of several cytoskeletal elements: myosin, ROCK, F-actin, RhoA, Cofilin and YAP/TAZ. Overexpression of the same cytoskeletal elements leads to inhibited patterning (C) and generally increased filopodia activity (D). All simulations were repeated 25 times. The mean and standard deviation (shaded colors) are shown.

In the case of over-expressed cytoskeletal elements (Fig. 6C-D), none of the simulated cases patterned beyond the original stiffness threshold; patterning was actually inhibited in most cases (Fig. 6C). This trend was accompanied by high filopodia activity throughout the whole range of simulated stiffnesses (Fig. 6D). This behavior for overexpression of cytoskeletal elements led to high nuclear YAP/TAZ for low stiffness levels onwards, low LFNG and DLL4 (and overall NICD) activity, in combination with high VEGFR2 expression, leading to decreased patterning and increased filopodia activity.

## Discussion

The processes underlying HHT2, involving a complex interplay of signaling pathways that collectively contribute to aberrant angiogenesis, remain incompletely characterized. This may have detrimental consequences for patients. In the present study, we specifically investigated the roles of YAP/TAZ stiffness mechanosensitivity and YAP/TAZ interplay with the NOTCH pathway in the context of angiogenesis dysregulated by ALK1 KO, known to cause HHT2. Upon ALK1 KO, our simulations showed increased nuclear YAP/TAZ and slow phenotypic pattern formation accompanied by high filopodia activity prior to pattern formation, indicating hypersprouting. As a consequence of YAP/TAZ stiffness sensitivity, these behaviors were mainly observed for low stiffnesses, whereas ALK1 KO cells more closely resembled wild-type behavior on higher stiffnesses. Moreover, the simulations indicated that ALK1 KO ECs were less likely to switch phenotype upon a positional shuffle compared to wild type, suggesting more migratory behavior. Furthermore, the simulations predicted that either direct or indirectly (via cytoskeletal elements) inhibition of YAP/TAZ could partially restore EC signaling and angiogenesis temporal dynamics, thus potentially attenuating the formation of aberrant vasculature due to disturbed phenotypic selection.

Previous computational studies have also used computational modelling to investigate EC behavior in HHT2, focusing on the EC polarization response to the mechanical regulator flow, and suggested junctional dysfunction as a new treatment target to prevent AVM formation(96). The critical role of temporal dynamics of signaling pathways has been gaining increasing attention, also in the field of pathogenesis, e.g. highlighting the critical role of LFNG, using in silico methods(29,42,43). In this study, we took those temporal dynamics into account and focused mainly on phenotypic selection, and the effect of stiffness, another important mechanical cue impacting YAP/TAZ signaling.

Our results are consistent with a recent experimental study showing that YAP/TAZ signaling may play a crucial role in HHT2(6). In their experiments, ALK1 KO led to pronounced upregulation of nuclear YAP/TAZ and to impaired endothelial polarization against the flow(6). This was restored by YAP/TAZ inhibition, which also prevented AVM formation(6). Since it has been shown that AVM formation is initiated by aberrant sprouting from veins(15), we hypothesized that targeting YAP/TAZ signaling due to its interactions with NOTCH, can also rescue the aberrant tip/stalk temporal dynamics caused by ALK1 KO. Our simulations predicted that ALK1 KO caused increased nuclear YAP/TAZ and VEGFR2 activity, and decreased LFNG expression and NOTCH activation, in agreement with experiments(6,25,26). The predicted hypersprouting is corroborated by observations in multiple experimental models(5,6,29). Therefore, the identified signaling interactions accurately describe and explain several experimental results focused on HHT.

In our simulations, inhibition of YAP/TAZ in ALK1 KO ECs led to faster patterning, in combination with lower filopodia, VEGFR2, and NOTCH activity. So, it partially normalized phenotypic selection of ALK1 KO ECs and potentially reduced hypersprouting, in spite of the protein levels not being completely restored to physiological signaling levels. This is consistent with our assumption that YAP/TAZ mainly affect LFNG and DLL4(39–41), while the ALK1 KO mutation mainly affects LFNG and HE. Targeting cytoskeletal elements, or other regulators of YAP/TAZ nuclearization(79), may help to prevent hypersprouting, but could also lead to off-target effects, for example affecting ROCK, which regulates vessel permeability(80). There should therefore be a careful trade-off between effectiveness of the chosen target and the potential off-target effects. Overall, our simulations indicate that the effects of YAP/TAZ, upregulated by the overactive VEGF-VEGFR2 signaling loop in ALK1 KO, play a major role in dysregulated phenotypic selection, that could be (partially) restored by inhibiting it.

Our simulations also highlight the importance of selecting multiple measurement timepoints in the experiments, given the oscillating nature of the signaling pathways involved. For example, VEGFR1 expression decreased only minimally upon ALK1 KO in our simulations, in contrast to experimental observations of Ola and Thalgott(5,25). A possible explanation could be a difference in the measured timepoint. In support of this, their data shows a dynamic temporal profile of VEGFR2 activity, with an initial steep increase that then gradually decreases. Our simulations indicated another important temporal component, showing initially decreased expression of HE, followed by increased expression, due to the positive VEGF feedback loop. This reinforces the notion that the dynamics of angiogenesis are characterized by temporal fluctuations that need to be taken into account in study design(42,51,57).

Environmental stiffness is another important mechanoregulator of angiogenic sprouting activity, in addition to flow mechanosensitivity(76). Our simulations indicated a shift in stiffness-responsive behavior of ALK1 KO ECs compared to wild-type ECs, with major effects in low stiffness regions specifically. This may explain why AVMs often occur in softer tissues, such as the liver, brain and lungs(2,4,8), all with stiffnesses generally below 6 kPa(46,47,73,81–83). Wild-type angiogenic sprouting has been shown to be optimal for intermediate stiffnesses, characterized by fast phenotypic patterning and dense vascular networks(42,43,83–85). Interestingly, experiments have shown that sprouting ECs contribute to a collection of rather heterogeneous stiffnesses in the sprout environment, potentially enhancing their own sprouting ability, as corroborated by our own previous simulations(41,86). This process might be dysregulated in HHT2 too; in fact, some ECs involved in vascular malformations secrete higher amounts of MMPs, leading to changes in stiffness and increased degradation of the extracellular environment thereby potentiating excessive sprouting(87,88). In addition to this, fluid shear stress has recently been suggested to play a major role in HHT2 through the mechanosensitive ion channel PIEZO1(89–92). Degradability, tissue stiffness and flow shear stress, are all mechanical cues shown to affect angiogenesis in HHT, further supporting the notion of mechanical stimuli as important contributors to the HHT2 pathology.

Cells with impaired LFNG signaling, such as ALK1 KO cells, showed increased positional shuffling in experiments(29), corroborated by the increased filopodia activity observed in our simulations. However, despite positional switching, our simulations indicated decreased phenotypical shuffling for ALK1 KO. Earlier studies have described and investigated this phenotypical shuffling and suggested the VEGFR1/VEGFR2 balance to be critical for a normal phenotypic switch(22). Our simulations predicted a disturbed balance with highly elevated VEGFR2 activation and minor changes to VEGFR1 activity, which could be one of the causes for the observed impaired shuffling, potentially through impairment of lateral inhibition. Phenotypic shuffling is required for physiological network formation, and previous experiments indicated that if phenotypic switching is reduced, this could lead to less branching (and potentially expansion instead of branching)(43). Hence, the decreased phenotypic switching capacity observed in the simulations indicates decreased phenotypic shuffling rates in ALK1 KO ECs, leading to less branching and instead vessel expansion, in agreement with vessel dilatation observed in HHT2(2,93). This observation of decreased phenotypic switching potential in ALK1 KO ECs is also supported by experiments indicating ectopic expression of arterial markers in veins that show signs of premature AVMs in mice(94). *In vitro*, VEGF needs to become inactive for ECs to achieve a venous phenotype(95). This might not be possible in ALK1 KO ECs, due to the overactive VEGF-VEGFR2-filopodia positive feedback loop. ALK1 KO ECs may have impaired differentiation capacity, preventing them from correctly forming all the endothelial subtypes necessary to obtain a healthy capillary bed. In later stages too, after tip/stalk pattern establishment, it was observed that LFNG-KO ECs did not return to their quiescent state(29). In summary, these findings suggest that ALK1 KO tip cells remain hyperactive, potentially unable to regulate NOTCH and VEGF activity, thereby unable to differentiate between the different phenotypes, leading to aberrant sprouting and erroneous arterio-venous connections. This could potentially be alleviated by YAP/TAZ inhibition, as the simulations indicated (partially) increased phenotypic switching potential in some settings.

In summary, the developed modelling framework has allowed us to investigate the phenotypic selection and shuffling behavior of ALK1 KO ECs over a range of stiffnesses, with and without YAP/TAZ inhibition. Our results highlight that ALK1 KO impairs phenotypic selection and shuffling by inhibiting NOTCH1 activation, especially in low stiffness environments, and indicate that targeting YAP/TAZ and endothelial stiffness sensitivity might provide promising therapeutic strategies to restore the physiological dynamics of angiogenesis.

## Methods

To elucidate the mechanisms underlying aberrant phenotypic selection by ECs in HHT2, we expanded our previously established modelling framework describing VEGF-NOTCH1 signaling and its crosstalk with YAP/TAZ in a 1D series of communicating ECs(41,97). More specifically, we extended the set of ordinary differential equations (ODEs) with the VEGF-FAK interaction, to allow for bidirectional feedback and split the generic VEGFR in the model into VEGFR1 and VEGFR2.

### Main model assumptions

#### Original modelling framework

In the modelled VEGF-NOTCH signaling cascade(97,98), simulated filopodia activity increases upon VEGF binding to VEGFR2, thereby increasing the VEGF sensing ability of the respective cell. At the same time, activated VEGFR2 upregulates DLL4 expression, leading to increased transactivation of NOTCH1 receptors on adjacent cells. Subsequent cleavage and translocation of the Notch1 Intracellular Domain (NICD) to the nucleus induces expression of target genes (HES/HEY, labelled here with HE) in those neighboring cells, which repress VEGFR2 expression, thereby decreasing their VEGF sensing abilities – a mechanism called lateral inhibition.

In Passier et al. we incorporated the mechanosensitivity of EC phenotypic selection by integrating the VEGF-NOTCH1 model with an ODE-based model describing YAP/TAZ nuclearization through cytoskeletal interactions(35,41). Briefly, substrate stiffness modulates the non-linear activity and clustering of adhesion proteins, generally represented and quantified in the model by the amount of phosphorylation of Focal Adhesion Kinase (FAK). Phosphorylated FAK (pFAK) subsequently activates RhoA. RhoA signaling promotes G-actin polymerization into F-actin via mDia, and stabilizes F-actin through ROCK mediated activation of LIMK, which inhibits the F-actin severing activity of Cofilin. Additionally, ROCK enhances myosin contractility, and together with F-actin, leads to the formation of contractile (actomyosin) stress fibers. These stress fibers are assumed to induce nuclear flattening – though not explicitly modelled -which enhances YAP/TAZ nuclear translocation(34,35). Importantly, the YAP/TAZ and NOTCH1 pathway are coupled via YAP/TAZ-mediated inhibition of both DLL4 and LFNG expression(39,40,99). This coupling enables mechanical cues, such as stiffness, to modulate VEGF-NOTCH1 signaling and endothelial phenotypic selection dynamics(41).

#### Extended model

Previous studies have included the effect of BMP9 by its upregulation of HE as well as LFNG(97). We used this to include ALK1, and to mimic the ALK1 KO mutation. We build on top of these studies by implementation of VEGF-FAK, to include the bidirectional crosstalk between the YAP/TAZ and VEGF-NOTCH pathways, which we hypothesize to be essential for both physiological as well as pathological angiogenesis, including that in HHT2 patients. Both stiffness as well as VEGF-dependent FAK activation rely on integrin interactions(100,101). At the same time, both enhance integrin activity too(102,103). Since integrins are not explicitly modeled in our framework, we assume a direct interaction between VEGF and FAK phosphorylation, that synergizes with the direct interaction between stiffness and FAK, thereby upregulating nuclear YAP/TAZ(38,68,69,104). Moreover, we adapted the parameters of the VEGFR parameter in the original model to reflect solely VEGFR2 behavior, and included VEGFR1, that acts as a decoy receptor, is generally much more abundantly present than VEGFR2(67) and is upregulated by HE(17,19,22). For all model interactions, see Fig. 7. In this section we highlight the modified and extended equations. For a complete overview of all model equations, please refer to appendix S1.

**Figure 7.**
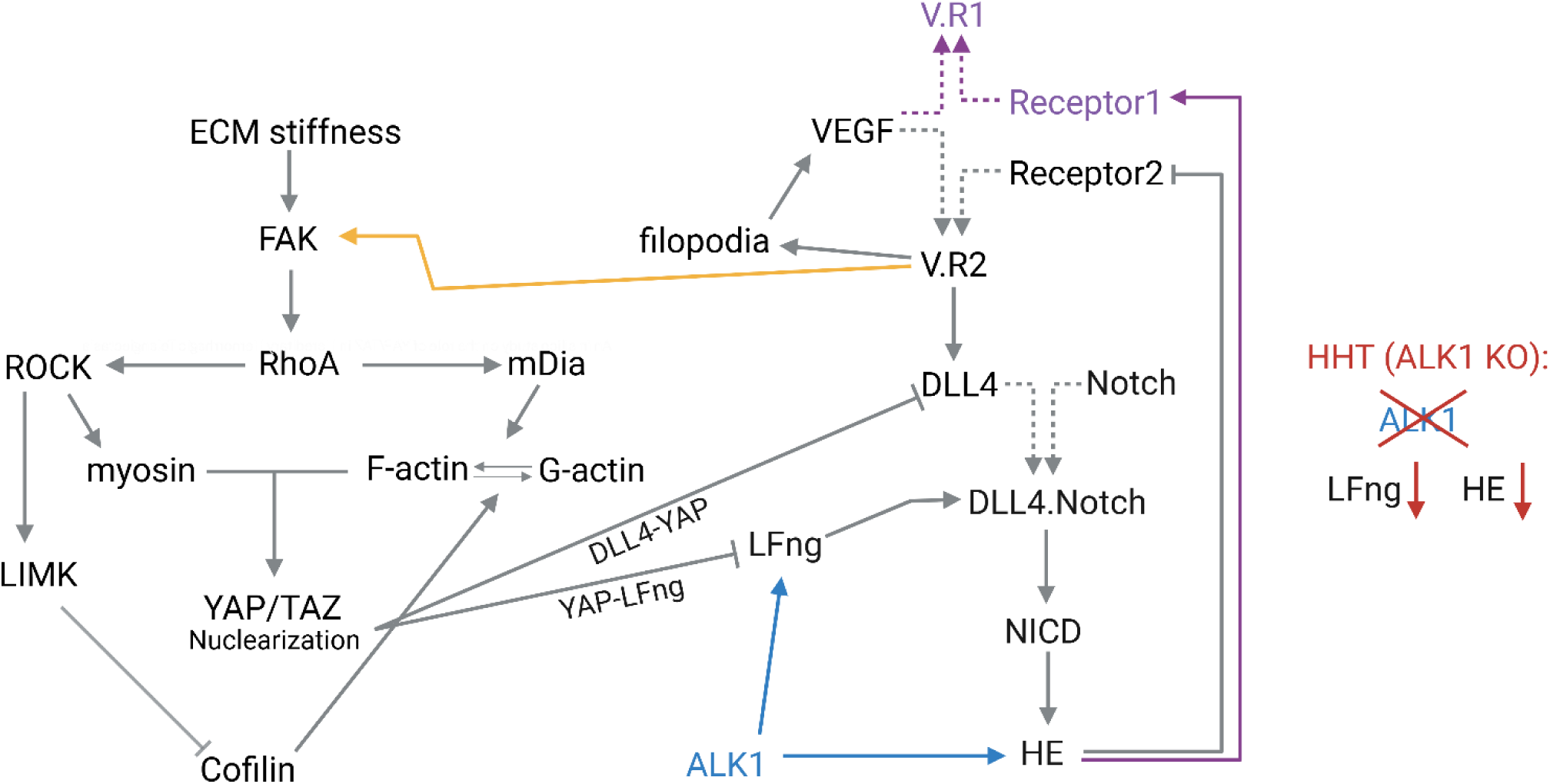
Schematic overview of the modelling framework. In gray all interactions present in the previous framework. In purple the interactions reflecting the split of VEGFR in VEGFR1 and VEGFR2, in yellow the interaction between VEGF and FAK, in blue the effects of ALK1 and in red those of ALK1 KO. Created using Biorender.com

### VEGF-mediated FAK phosphorylation

The original equation for FAK (*φ*) phosphorylation(35,41)was expanded to include active VEGFR2 signaling (*V*. *R*_2_) as an additional modulator that synergizes with the effects of stiffness (*E*):

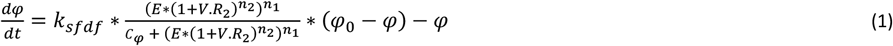

in which *φ*_0_ denotes the total amount of FAK, *k*_*sfdf*_ represents the net rate of FAK activation and dephosphorylation and *C*_*φ*_ acts as a Michaelis-Menten-like constant, indicating the inflection point where there is a shift in behavior resulting from the combined effect of the pro-adhesion formation stimuli – stiffness (*E*) and activated VEGFR2 (*V*. *R*_2_) – for which the phosphorylation rate of FAK reaches half of its maximum. *V*. *R*_2_ was introduced in a 1 + *V*. *R*_2_ formulation, to ensure that stiffness can still induce YAP/TAZ nuclearization independently of VEGF(105). Moreover, a distinct exponent (*n*_2_) was applied to the *V*. *R*_2_ term to capture the experimentally observed non-linear dependence of phosphorylated FAK on VEGF(104).

As a result of this implemented feedback mechanism, nuclear YAP/TAZ levels can become heterogeneous across the row of ECs, in contrast to their previous homogeneous distribution. Given that nuclear YAP/TAZ modulates the binding affinity between DLL4 and NOTCH1, this introduces heterogeneity with respect to binding affinities too. Moreover, to accurately reflect interactions on both cell edges, we distinguish between the left and right part of each modelled EC, allowing for spatially resolved NOTCH1-DLL4 interactions on both edges. Consequently, cell *i* interacts with cell *i* − 1 on its left side and with cell *i* + 1 on its right side. Consistent with the previously described mechanism of Fringe-mediated glycosylation of the NOTCH1 receptor determining the binding affinity(27,28), we assign the binding affinity 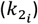 of the NOTCH1 receptor-bearing cell. Accordingly, when DLL4 on the left side of cell *i* interacts with NOTCH1 on the right side of cell *i* − 1, the governing binding affinity is that of cell *i* − 1. An example below:

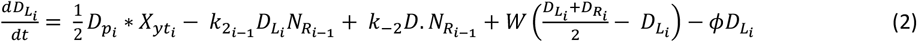

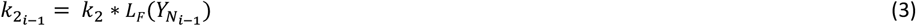

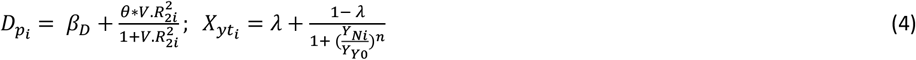

where *D*_*p*_ represents the production of DLL4, determined by the basal production rate (*β*_*D*_) and the stimulatory effect of activated *V*. *R*_2_, scaled by *θ*. This term is multiplied with 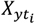 to account for the repressing effect of YAP/TAZ, governed by the amount of nuclear YAP/TAZ 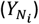, a baseline value of nuclear YAP/TAZ (*Y*_*Y*0_), a scaling factor (*n*) and *λ*, determining the maximum effect of this term (e.g. an *λ* of 0.5 indicates a maximum repressing effect of 50%). 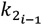 indicates the binding affinity of DLL4 on the left side of cell *i* to bind with NOTCH on the right side of cell *i* – 1 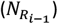, as determined by the amount of nuclear YAP/TAZ in that cell 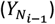, a basal value for the binding affinity (*k*_2_) and LFNG, linearly taken into account by *L*_*F*_. Bound DLL4-NOTCH 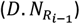 can also dissociate, with probability *k*_−2_. Additionally, *W* represents diffusion of DLL4 among the left and right cell edges and *ϕ* indicates degradation of DLL4.

### Implementation of the differential roles of VEGFR1 and VEGFR2

In addition to incorporating the VEGF-FAK interaction, we extended the model by differentiating between VEGFR2 and VEGFR1. To account for competition of VEGF-receptor binding, reflecting the spatial and availability constraints, we introduced the ratio (*ξ*) between VEGFR1 (*R*_1_) and VEGFR2 (*R*_2_), representing the generally relative abundance of *R*_1_ compared to *R*_2_(65,67). This ratio is incorporated as a scaling factor modulating the effective binding rate (*k*_1_) of VEGF to its respective receptors. This gives rise to the following formulas:

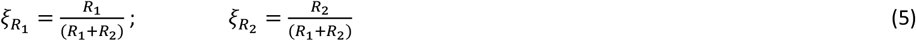

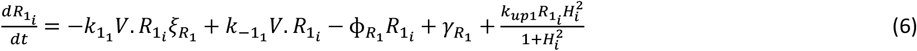

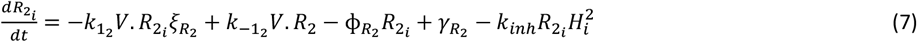

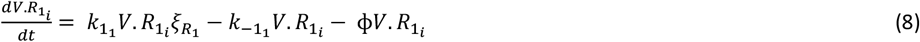

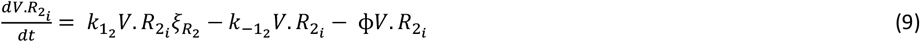

in which *k*_−1_ denotes the dissociation rate of the VEGF-Receptor complex (*V*. *R*), ɸ represents the degradation rate and *γ* indicates the basal production rate. By distinguishing between *R*_1_and *R*_2_, we can explicitly implement the different regulatory effects of NOTCH target genes (*H*) on the different receptors. More precisely, *H* upregulates *R*_1_ expression (via *k*_*up*1_), which is implemented analogously to the downregulatory effect of *H* on *R*_2_ (via *k*_*inh*_). To prevent excessive accumulation and numerical instability of unbound *R*_1_, the upregulatory term is divided by an additional *H*-dependent denominator, analogous to the formulation used for DLL4 upregulation upon *V*. *R* activation.

### Mimicking ALK1 KO

In correspondence with previous work(97), we implemented the effects of BMP9 signaling through ALK1, by implementation of a parameter affecting both LFNG, and through that, the binding affinity 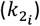, as well as the HE basal production rate (*β*_*HE*_). In case of ALK1 KO (HHT2), this parameter (*B*_*eff*_), decreases to below one to lower LFNG and HE activity, while in case of BMP9 stimulation, this parameter is set to a value above one:

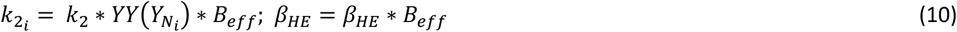

### Parameter choices & fitting

Most of the parameters already present in the model were retained from previous studies(35,41,97,98). However, specific parameters related to FAK phosphorylation (see equation 1) were adjusted to accommodate the addition of the VEGF-FAK interaction, while maintaining consistency with previously captured and validated model behaviors. More specifically, the net activation/dephosphorylation rate (*k*_*sfdf*_), Michaelis-Menten constant (*C*) and the Hill exponent governing the overall nonlinearity of FAK phosphorylation with respect to stiffness (*n*_1_) were recalibrated. These values, as well as the exponent specifically for the (1 + *V*. *R*_2_) term (*n*_2_) were fitted to experimental data obtained by Abedi et al, who observed increasing FAK phosphorylation with increasing VEGF concentrations(104). Initial parameter estimation was conducted using the least-squares non linear fitting algorithm of MATLAB (lsqnonlin), which provided approximate parameter ranges that minimized the discrepancy between the model output and experimentally observed FAK phosphorylation for increasing VEGF levels. However, in addition to reproducing this data, the model output needed to satisfy a list of other qualitative and quantitative constraints. Specifically, nuclear YAP/TAZ should exhibit a steep increase for lower stiffness regimes, and continue to increase, more gradually, for higher stiffness levels(70). Moreover, the model needed to reproduce the biphasic response with respect to increasing stiffnesses and should cease patterning around 24 kPa, consistent with original model behavior(41). In addition to this, VEGF concentrations slightly above the levels used to instigate physiological patterning, should yield a further increase in both phosphorylated FAK and nuclear YAP/TAZ, avoiding saturation at physiological VEGF levels. To address these additional criteria, final parameter tuning was performed manually, using the initial fitted values as starting points.

In addition to the adjustment of FAK-phosphorylation parameters, several new parameters were introduced to account for the split of the VEGF receptor into *R*_1_and *R*_2_. These parameters were mainly based on previous experimental studies, indicating higher binding affinity between *R*_1_ and VEGF compared to *R*_2_(106), increased degradation rates for *R*_1_ compared to *R*_2_, based on their respective half life times(107) and a larger effect of transcriptional regulation by *H* on *R*_1_ than on *R*_2_(22). The degradation and dissociation rates of the bound *V*. *R*_1_ complex were assumed to be equivalent to those of *V*. *R*_2_. To estimate the relative production rate of *R*_1_, we used available data reporting average ratios between *R*_1_ and *R*_2_ allowing us to calibrate baseline production of *R*_1_ accordingly(65,67). Lastly, we increased the baseline level of VEGF, to ensure the possibility of patterning in the presence of *R*_1_, without disrupting previously established model dynamics(41).

To determine the scaling factors required to mimic the effects of ALK1 KO on the production of *H* and LFNG activity, we used qPCR data of Ristori et al(97). Their study reported a significant decrease of NOTCH target genes HES/HEY (*H*) expression in ALK1 KO cells. Please refer to appendix S2 for a complete overview of all parameter values.

### Model validation

In addition to fitting the model parameters on a wide range of available experimental data to ensure biologically accurate behavior, we performed model validation simulations against independent experimental observations, for several different scenario’s. Following each model alteration, we verified that essential behaviors captured in the original model were preserved(41). One of the main aspects we considered was nuclearization of YAP/TAZ in response to stiffness. Previous experimental studies have demonstrated that nuclear YAP/TAZ levels rise with increasing stiffness, showing an initial, sharp increase for the low stiffness regime and a more gradual increase for higher ranges of stiffness (70 kPa and up)(70). Moreover, an essential element of the angiogenic mechanoresponse that we captured with our previous *in silico* model, is the existence of an optimal stiffness, yielding fast and robust patterning(41,84,85). Throughout all steps, we verified that the model retained these essential mechanobiological features.

For the pathological scenario simulated in this study, HHT2 (simulated via ALK1 KO), we employed distinct experimental data to validate, as the HHT2 scenario deviates from the original, physiological model context. To this end, model output was compared with data of Park et al, who reported elevated YAP/TAZ nuclearization upon ALK1 KO and demonstrated that pharmacological inhibition of YAP/TAZ prevented AVM formation in ALK1 KO mice(6). Additionally, VEGF receptor dynamics were validated against data of Ola et al, who showed that loss of ALK1 results in increased activity of VEGFR2, in spite of unchanged overall VEGFR2 levels, in combination with decreased levels of VEGFR1(25).

### Model simulations

Following validation of the extended modelling framework, we performed a series of simulations to explore EC behavior in different settings. Firstly, we simulated ALK1 KO and analyzed how behavior of the different angiogenic signaling elements was affected. Subsequently, we simulated pharmacological inhibition of YAP/TAZ, in line with experiments conducted by Park et al(6), to assess to what extent the effect of ALK1 KO on those key signaling elements could be mitigated by YAP/TAZ inhibition. To further investigate endothelial plasticity, we examined the tendency of cells to undergo a phenotypic switch, in response to switching the environmental cue, comparing behavior between wild-type and ALK1 KO ECs. In addition to YAP/TAZ KO, we explored alternative intervention targets for their potential of restoring patterning and shuffling behavior upon knock down or overexpression. Moreover, we investigated the role of extracellular matrix stiffness under ALK1-deficient conditions, both in the absence and presence of interventions aimed at rescuing patterning.

All simulations were conducted using MATLAB 2024a, employing the ODEsolver ode15s. Initial conditions were set to zero for all intracellular proteins, and unless specified otherwise, simulations started with a 24h acclimatization period (the absence of external cues), after which specific stimuli, such as VEGF, were introduced, mimicking experimental protocols. To account for inherent biological VEGF sensing discrepancies between cells (e.g. due to differences in cell shape or position), a small random factor (10^−16^) was added to the VEGF input. Baseline stiffness was set to 5 kPa, to reflect physiological liver stiffness(46,47,73). Lastly, periodic boundary conditions were imposed, to enable interaction between the first and last cell of the row, so maintaining continuity in the simulated environment.

#### Cell shuffling

The process of phenotypic shuffling was simulated by switching to a two-cell model. Periodic boundary conditions were removed, limiting cell-cell interactions to a single interface (the right edge of cell 1 and the left edge of cell 2). As a consequence, all produced DLL4 and NOTCH1 was restricted to this one edge and intracellular diffusion over the edges was removed. The simulations were divided into two sequential 24 hour-phases: (i) exposure to VEGF, with a slight bias towards one cell (0.9*VEGF and 1.0*VEGF), to bias tip cell selection; (ii) a gradual transition (over 2 hours(22)) towards a new asymmetric VEGF distribution (see equation 11), favoring the tip phenotype in the opposite cell. The magnitude of the asymmetric VEGF cue (*α*_*s*_) used in phase (ii) of these simulations was systematically varied, ranging from 10 to 90% (*α*_*s*_ being between 0.1 and 0.9 respectively) for cell 2 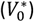 and 100% of the original VEGF cues (*V*_0_) for cell 1 respectively, to examine its effects on phenotypic shuffling in both physiological and ALK1 KO conditions (see equation 11). In addition to this asymmetric VEGF cue, the cells were exposed to an asymmetric DLL4 cue in phase ii, representing DLL4 levels of a neighboring tip and stalk cell respectively (as if the cells had switched position and acquired new neighbors).

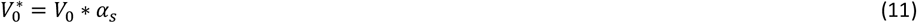

#### Intervention targets

To mimic knock down of possible intervention targets, we multiplied the (de)activation rates of the respective elements with a factor 10, consistent with previous work(35,41). More specifically, the deactivation rates of Myosin (*k*_*dmy*_), ROCK (*k*_*dROCK*_), RhoA (*k*_*dp*_) and Cofilin (*k*_*cr*_) were multiplied by 10, mimicking Blebbistatin, Y-27362, Rhosin and SZ-3(108–111), while the F-actin polymerization rate (*k*_*ra*_) was decreased with a factor 10 to simulate latrunculin(112). We mimicked verteporfin, which inhibits YAP/TAZ by preventing its translocation to the nucleus and blocking YAP-TEAD formation, by setting the YAP/TAZ active nuclear translocation rate to zero, only allowing for basal nuclearization activity(6,113–115). It should be noted, that in the model we do not explicitly distinguish between different ways of inhibiting or knocking out YAP/TAZ, which is why YAP/TAZ inhibition is referred to as ‘YAP/TAZ KO’. For completeness, we additionally simulated cytoskeletal overexpression by dividing the respective deactivation rates by 10, or by multiplying with 10 in case of the polymerization rate for F-actin. For all simulations, cells were exposed to the standard, symmetric, VEGF cue to instigate patterning.

#### External cues

The effect of extracellular matrix stiffness was investigated by performing the simulations mentioned above across a broad stiffness range, from 0.5 to 250 kPa. The step size was incrementally increased for higher stiffnesses, to reflect decreasing sensitivity of YAP/TAZ nuclearization to stiffness. Beyond 250 kPa, no substantial changes in model outputs were observed.

Across all simulations, phenotypic selection time was defined and quantified consistently. ECs were classified as tip cells when their filopodia activity surpassed the threshold set at 20, while stalk cells needed to have a filopodia activity lower than this threshold, in accordance with previous studies(56,97). A pattern was considered established when at least 40% of the ECs adopted the tip cell phenotype, without any of those tip cells being adjacent. To account for variability, all simulations were repeated 25 times. Moreover, average activity and expression of all outputs of interest were determined by averaging their amounts over the timepoints before patterning occurred.

## Data availability statement

All data and code will be available at 4TU. ResearchData, through: https://doi.org/10.4121/b871982a-763e-4ff4-8b0e-3e01f77799ec

## Acknowledgements

The authors acknowledge Shayan Zarin-Bal for the insightful discussions. The schematics depicted in figures 2, 3, 4 and 7 were created with Biorender.com.

## Author contributions

Conceptualization (TR, MP), Formal Analysis (MP), Funding Acquisition (TR), methodology (TR, MP), supervision (TR, SL), visualization (MP), writing - original draft (MP), writing - review & editing (TR, SL, MP).

