## Supplementary Information for "Targeting stiffness-dependent YAP/TAZ restores angiogenesis dynamics impaired by ALK1 knockout *in silico*"

**Supplementary Information 1**

This section provides the model equations including explanation.

Our previous iteration of the model, describing the interactions between VEGF-NOTCH and YAP/TAZ signaling in a row of communicating endothelial cells(1) was extended to include the VEGF-FAK interaction as well as distinctive VEGFR1 and VEGFR2 receptors (instead of one generic VEGF receptor). The row of ECs was simulated using periodic boundary conditions, while we distinguished between the left (subscript ‘L’) and right (subscript ‘R’) cell edge. To indicate a specific cell in the row of ECs, we used the subscript ‘i’, to indicate the i^th^ cell, with i-1 indicating the left neighbor of this cell and i+1 the right neighbor. Below, we provide an overview of the full model and the equations it comprises.

**VEGF-NOTCH signaling**

We considered the same assumptions as in the original version of the VEGF-NOTCH1 signaling framework, developed by Venkatraman et al(2), and extended by Ristori et al(3). Hence, the amount of sensed VEGF ($V_{i}$) depends on the amount of VEGF ($V_{0}$) that the cells are continuously exposed to, in addition to the positive feedback provided by filopodia on that cell ($f_{i}$):

$V_{i}=V_{0}+k_{3}V_{0}f_{i}^{2}$	(S1)

which is scaled by parameter $k_{3}$. VEGF can then bind to the either one of the VEGF receptors ($R_{1_{i}}$ or $R_{2_{i}}$), thereby establishing a VEGF-VEGFR bound complex ($V.R_{1_{i}}$ and $V.R_{2_{i}}$), leading to upregulation of filopodia formation in case of activated $R_{2}$. To reflect the spatial and availability constraints of VEGF binding to the different VEGF receptors, we introduced the ratio ($\xi$) between the different receptors. This represents the relative abundance of $R_{1}$ compared to $R_{2}$. This ratio scales the effective binding rate ($k_{1}$) of VEGF to the respective receptors. The amount of VEGFR available for binding with VEGF is downregulated ($k_{inh}$) by Hes1/Hey1 ($H_{i}$) in case of $R_{2}$, while it is upregulated ($k_{up1}$) in case of $R_{1}$, giving rise to the following equations:

$\xi_{R_{1}}=\frac{R_{1}}{(R_{1}+R_{2})}$ ;	$\xi_{R_{2}}=\frac{R_{2}}{(R_{1}+R_{2})}$	(S2)

$\frac{dR_{1_{i}}}{dt}={-k}_{1_{1}}VR_{1_{i}}\xi_{R_{1}}+k_{{-1}_{1}}V.R_{1_{i}}-ɸ_{R_{1}}R_{1_{i}}+\gamma_{R_{1}}+\frac{k_{up1}R_{1_{i}}H_{i}^{2}}{1+H_{i}^{2}}$	(S3)

$\frac{dR_{2_{i}}}{dt}={-k}_{1_{2}}VR_{2_{i}}\xi_{R_{2}}+k_{{-1}_{2}}V.R_{2}-ɸ_{R_{2}}R_{2_{i}}+\gamma_{R_{2}}-k_{inh}R_{2_{i}}H_{i}^{2}$	(S4)

$\frac{d{V.R}_{1_{i}}}{dt}= k_{1_{1}}{VR}_{1_{i}}\xi_{R_{1}}-k_{{-1}_{1}}V.R_{1_{i}}- ɸ{V.R}_{1_{i}}$	(S5)

$\frac{d{V.R}_{2_{i}}}{dt}= k_{1_{2}}{VR}_{2_{i}}\xi_{R_{2}}-k_{{-1}_{2}}V.R_{2_{i}}- ɸ{V.R}_{2_{i}}$	(S6)

$\frac{{df}_{i}}{dt}= \beta+ k_{f}{V.R}_{2_{i}}^{2}- k_{-f}f_{i}$	(S7)

where $k_{-1}$ represents the dissociation rate of the bound VEGF-receptor complex ($V.R$), $\gamma$ represents the basal production rate of the respective receptors and $ɸ$ indicates the degradation rate. $k_{-f}$ and $k_{f}$ are filopodia specific, and indicate the filopodia turnover rate and formation rate affecting the upregulation by $V.R_{2_{i}}$ respectively. $\beta$ indicates basal formation of filopodia. Upon activation of $R_{2}$, DLL4 ($D_{i}$) gets upregulated, allowing binding with the NOTCH1 receptor ($N_{i}$), leading to complex formation (${D.N}_{i}$) and subsequent activation of NOTCH1. Additionally, external DLL4 ($D_{ext}$) can bind to NOTCH1, leading to similar complex formation ($D_{ext}.N_{i}$) and subsequent activation. This gives rise to the following equations, shown here for the left cell edge, but this is analogous for the other edge:

$\frac{{dD}_{L_{i}}}{dt}=\frac{1}{2}(\beta_{D}+\frac{\theta*V.R_{2i}^{2}}{1+V.R_{2i}^{2}}*X_{{yt}_{i})}- k_{2_{i-1}}D_{L_{i}}N_{R_{i-1}}+ k_{-2}D.N_{R_{i-1}}+W\left( \frac{D_{L_{i}}+D_{R_{i}}}{2}- D_{L_{i}} \right)-\phi D_{L_{i}}$	(S8)

$\frac{{dN}_{L_{i}}}{dt}=\frac{\gamma}{2}- k_{2_{i}}N_{L_{i}}{(D}_{R_{i-1}}+D_{ext})+k_{-2}({D.N}_{L_{i}}+D_{ext}.N_{L_{i}})+W(\frac{N_{L_{i}}+ N_{R_{i}}}{2}- N_{L_{i}})-\phi N_{L_{i}}$	(S9)

$\frac{{D.N}_{L_{i}}}{dt}=k_{2_{i}}N_{L_{i}}D_{R_{i-1}}-k_{-2}{D.N}_{L_{i}}-k_{cat}{D.N}_{L_{i}}-\phi{D.N}_{L_{i}}$	(S10)

$\frac{D_{ext}.N_{L_{i}}}{dt}= k_{2_{i}}N_{L_{i}}D_{ext}-k_{-2}D_{ext}.N_{L_{i}}-k_{cat}D_{ext}.N_{L_{i}}- \phi D_{ext}.N_{L_{i}}$	(S11)

in which θ scales the downregulation of DLL4 production by ${V.R}_{i}$, $\beta$ indicates basal production, $W$ indicates the diffusion of unbound DLL4 and NOTCH over the cell edges, $k_{cat}$ the catalysis of the ${D.N}_{i}$ complex and $k_{2_{i}}$ and $k_{-2}$ the association and dissociation rates of the complex respectively. Due to the bidirectional feedback between the YAP/TAZ and NOTCH signaling pathway, YAP/TAZ levels can become heterogeneous across the ECs. Since YAP/TAZ modulates the binding affinity of DLL4 and NOTCH via LFng, the binding affinity ($k_{2_{i}}$) also becomes heterogeneous across cells. To account for the glycosylation effect by LFng on the NOTCH1 receptor, we assign the binding affinity for the DLL4-NOTCH1 interactions of the NOTCH1 bearing cell. As an example; if DLL4 on the left cell edge of cell i binds to NOTCH1 on the right edge of cell i-1, the governing binding affinity is that of cell i-1, as can be seen in S8-10. The effect of LFng on the binding affinity was taken into account linearly, via $a$ and $b$ :

$k_{2_{i-1}}= k_{2}*\left( a*Y_{N_{i-1}}+b \right)$	(S12)

where $k_{2}$ represents the original value for the binding affinity. The inhibitory interaction between YAP/TAZ and DLL4, represented in the formulas by $X_{yt}$, was modelled using the following equation:

$X_{{yt}_{i}}= \left( \lambda+\frac{1-\lambda}{1+\left( \frac{Y_{Ni}}{Y_{Y0}} \right)^{n}} \right)$	(S13)

in which $\lambda$ represents the maximum inhibitory effect, $Y_{Y0}$ the threshold level of nuclear YAP/TAZ from which onwards there should be an inhibitory effect, $Y_{Ni}$ indicating the nuclear YAP/TAZ fraction and $n$ representing the scaling factor. Finally, activated NOTCH1 is cleaved, resulting in the NOTCH1 Intracellular Domain (NICD) ($I_{i}$), which in turn leads to increased expression of Hes1/Hey1, via:

$\frac{{dI}_{i}}{dt}=k_{cat}\left( {D.N}_{L_{i}}+{D.N}_{R_{i}}+ D_{ext}.N_{L_{i}}+ D_{ext}.N_{R_{i}} \right)-\phi I_{i}$	(S14)

$\frac{dH_{i}}{dt}= \beta_{HE}^{*}+\theta\frac{I_{i}^{2}}{1+I_{i}^{2}}- \phi H_{i}$	(S15)

**YAP/TAZ signaling**

We additionally adapted our previous YAP/TAZ signaling part of the model, which was based on the model developed by Sun et al(4). In the model, initial stiffness sensation happens through focal adhesion kinase, FAK ($\varphi$), which can be phosphorylated ($\varphi_{p}$) resulting from integrin and protein clustering. This is modelled via Michaelis-Menten kinetics. The parameters in this equation were adapted to accurately represent EC behavior, taking into account that not only stiffness ($E$) affects FAK phosphorylation, but active VEGFR2 signaling (${V.R}_{2}$) too, leading to:

$\frac{d\varphi_{p_{i}}}{dt}=k_{sfdf}*\frac{\left( E*\left( 1+{V.R}_{2_{i}} \right)^{n_{2}} \right)^{n_{1}}}{{C_{\varphi} + \left( E*\left( 1+{V.R}_{2_{i}} \right)^{n_{2}} \right)}^{n_{1}}}*\left( \varphi_{0}-\varphi_{i} \right)-\varphi_{i}$	(S16)

in which $\varphi_{0}$ represents the total amount of FAK present, $k_{sfdf}$ denotes the net FAK activation/dephosphorylation and $C_{\varphi}$ represents the Michaelis-Menten constant, providing an inflection point at which the behavior shifts resulting from the combined effect of pro-adhesion formation stimuli - activated VEGFR2 and stiffness – for which the phosphorylation rate of FAK is at half of its maximum value. Finally, $n_{1}$ and $n_{2}$ represent distinctive factors scaling the different effects on FAK phosphorylation. Activation of such adhesion molecules allows RhoA ($\rho$) to bind GTP, leading to actomyosin formation through mDia ($\sigma$) and ROCK ($\omega$) activation, via:

$\frac{d\rho_{i}}{dt}= k_{fkp}\left( \upsilon\varphi_{p_{i}}^{2}+1 \right)\left( \rho_{0}-\rho_{i} \right)-k_{dp}\rho_{i}$	(S17)

$\frac{d\sigma_{i}}{dt}=k_{mp}\rho_{i}\left( \sigma_{0}-\sigma_{i} \right)-k_{dmdia}\sigma_{i}$	(S18)

$\frac{d\omega_{i}}{dt}=k_{rp}\rho_{i}\left( \omega_{0}-\omega_{i} \right)-k_{drock}\omega_{i}$	(S19)

where, similar to the total amount of FAK, $\rho_{0}$*,* $\sigma_{0}$ and $\omega_{0}$ represent the total amounts of RhoA, mDia and ROCK respectively. Baseline activation of RhoA by FAK is indicated by $\upsilon$, activation by other mechanisms by $k_{fkp}$ and the deactivation rate is represented by $k_{dp}$. The activation rates of mDia and ROCK by FAK are represented by $k_{mp}$ and $k_{rp}$, whereas the respective deactivation rates are represented by $k_{dmdia}$ and $k_{drock}$. Upon activation of ROCK, LIM kinase, LIMK ($L$) as well as myosin ($M$) activity are promoted by kinase phosphorylation and inhibition of phosphatase, via:

$\frac{dM_{i}}{dt}=k_{mr}(\varepsilon T+1)\left( M_{0}-M_{i} \right)-k_{dmy}M_{i}$	(S20)

$\frac{dL_{i}}{dt}=k_{lr}\left( \tau T+1 \right)\left( L_{0}-L_{i} \right)-k_{dl}L_{i}$	(S21)

for which $M_{0}$ and $L_{0}$ indicate the total amounts of Myosin and LIMK, $k_{mr}$ and $k_{lr}$ denote the activation rates by pathways other than ROCK and $k_{dmy}$ and $k_{dl}$*_­_* indicate the deactivation rates. Additionally, myosin as well as LIMK are activated and amplified by ROCK, scaled with $\varepsilon$ and $\tau$. This interaction is incorporated using a smoothing function ($T$), so mimicking the requirement that ROCK activity needs to exceed a threshold in order to activate LIMK and Myosin. This was implemented using the built-in error function ($erf$) in MATLAB, through:

$T=\frac{e^{-S_{\varphi}^{2}(\varphi_{i}-\varphi_{S})^{2}}}{2 \sqrt{\pi}S_{\varphi}}+\frac{\varphi-\varphi_{S}}{2}(1+erf(S_{\varphi}(\varphi_{i}-\varphi_{S}))$	(S22)

in which the activator (ROCK) has a specific activation threshold ($\varphi_{S}$), as well as a parameter ($S_{\varphi}$) determining the sharpness of the response. Consecutively, LIMK can inactivate Cofilin ($C$), which normally severs F-actin ($F$) through phosphorylation. Additionally, mDia polymerizes G-actin to F-actin, as represented by:

$\frac{dC_{i}}{dt}=k_{to}\left( C_{0}-C_{i} \right)-k_{cr}(1-k_{ll}l_{0}){L_{i}}^{2}C_{i}$	(S23)

$\frac{dF_{i}}{dt}=k_{ra}\left( \alpha T+1 \right)(F_{0}-F_{i})-k_{dep}F_{i}-k_{fc1}C_{i}F_{i}$	(S24)

in which $C_{0}$ and $F_{0}$ indicate the total amounts of Cofilin and F-actin respectively. $k_{cr}$ represents the phosphorylation rate of cofilin due to LIMK, while $k_{to}$ and $k_{ll}$ represent the general dephosphorylation and phosphorylation inhibition rate caused by the combination of LIMK and LATS ($l_{0}$) respectively. For F-actin, $k_{dep}$ represents the general depolymerization rate, while $k_{fc1}$ represents the disassembly rate due to Cofilin, and $k_{ra}$ indicates the polymerization rate. The amplification of polymerization caused by mDia is represented by $\alpha$, and is incorporated using the smoothing function, $T$. Finally, stress fibers are formed by activated myosin and F-actin, causing nuclear flattening, allowing nuclear translocation of YAP/TAZ ($Y_{N}$), described by:

$\frac{dY_{N_{i}}}{dt}=\left( k_{cn}+k_{cy}F_{i}M_{i} \right)\left( Y_{0}-Y_{N_{i}} \right)-(k_{nc}+k_{ly}l_{p})Y_{N_{i}}$	(S25)

in which basal YAP/TAZ nuclear translocation and YAP/TAZ nuclear translocation caused by stress fibers are described by $k_{cn}$ and $k_{cy}$ respectively. Translocation from the nucleus to the cytoplasm with and without active LATS ($l_{p}$) is described by $k_{ly}$ and $k_{nc}$. Finally, total YAP/TAZ is described by $Y_{0}$.

**Mimicking ALK1 KO**

To incorporate the effects of ALK1 KO, we included a factor ($B_{eff}$), that scales the binding affinity ($k_{2_{i}}$) and basal production rate of Hes1/Hey1 ($\beta_{HE}$), in the following manner:

$k_{2_{i}}^{*}= k_{2}*\left( a*Y_{N_{i}}+b \right) *B_{eff1};$	$\beta_{HE}^{*}=\beta_{HE}*B_{eff2}$	(S26)

**Supplementary information 2**

This section contains all parameter values used to obtain the data as shown in the paper.

| **Param.** | **Value** | **Description** |
| --- | --- | --- |
| $V_{0}$ | 0.23 / 0.023 [cu] | Reference amount of VEGF, depending on the scenario |
| $k_{3}$ | 0.005 [cu^-2^] | Factor scaling positive feedback between filopodia and VEGF |
| $k_{1_{2}}$ | 0.1 [cu^-1^ sec^-1^] | Rate of association of V.R2 |
| $k_{-1_{2}}$ | 0.001 [sec^-1^] | Rate of dissociation of V.R2 |
| $\phi$ | 0.005 [sec^-1^] | Protein degradation |
| $\gamma$ | 0.005 [cu sec^-1^] | Protein production |
| $k_{inh}$ | 0.003 [cu^-2^ sec^-1^] | Scales the impact of inhibition of VR2 by H |
| $\beta$ | 0.001 [cu sec^-1^] | Basal filopodia and Hes1/Hey1 formation |
| $k_{f}$ | 0.1 [sec^-1^] | Filopodia formation rate |
| $k_{-f}$ | 0.001 [sec^-1^] | Rate of filopodia turnover |
| $\theta$ | 0.1[sec^-1^] | Downregulation rate of DLL4 production by V.R |
| $k_{-2}$ | 0.1 [sec^-1^] | Dissociation rate of D.N |
| $W$ | 0.001 [sec^-1^] | Diffusion of unbound Notch and DLL4 across both cell edges |
| $k_{cat}$ | 0.1 [sec^-1^] | Catalysis rate of D.N |
| $k_{2}$ | 0.002 [cu^-1^ sec^-1^] | Association rate of D and N |
| $k_{sfdf}$ | 0.194 [] | Net activation/dephosphorylation rate of K |
| $\upsilon$ | 500 [] | Activation rate of RhoA, by FAK |
| $k_{fkp}$ | 0.018 [sec^-1^] | Activation rate of RhoA through other mechanisms than FAK |
| $k_{dp}$ | 0.625 [sec^-1^] | Deactivation rate |
| $k_{rp}$ | 2.2 [sec^-1^] | Activation rate of ROCK by RhoA |
| $k_{mp}$ | 1 [sec^-1^] | Activation rate of mDia by RhoA |
| $k_{drock}$ | 0.8 [sec^-1^] | Deactivation rate of ROCK |
| $k_{dmdia}$ | 1 [sec^-1^] | Deactivation rate of mDia |
| $k_{mr}$ | 0.015 [sec^-1^] | Activation of Myo by other pathways than ROCK |
| $k_{lr}$ | 0.07 [sec^-1^] | Activation of LIMK by other pathways than ROCK |
| $k_{dmy}$ | 0.067 [sec^-1^] | Deactivation of Myo |
| $k_{dl}$ | 2 [sec^-1^] | Deactivation of LIMK |
| $\varepsilon$ | 40 [] | Activation of Myo through ROCK |
| $\tau$ | 200 [] | Activation of LIMK through ROCK |
| $S_{\varphi}$ | 13 [] | Smoothing parameter, determining sharpness of the activation – ROCK |
| $\varphi_{s}$ | 0.26 [] | Activation threshold for the smoothing function - ROCK |
| $S_{\sigma}$ | 10 [] | Smoothing parameter, determining sharpness of the activation – mDia |
| $\sigma_{S}$ | 0.13 [] | Activation threshold for the smoothing function - mDia |
| $k_{to}$ | 0.04 [sec^-1^] | Dephosphorylation rate of cofilin |
| $k_{cr}$ | 0.7 [sec^-1^] | Rate of phosphorylation of cofilin by LIMK |
| $k_{ll}$ | 0.8 [sec^-1^] | Phosphorylation inhibition rate of cofilin by LIMK, resulting from LATS_0_ |
| $k_{ra}$ | 0.4 [sec^-1^] | Polymerization rate of cytoplasmic F-actin |
| $k_{dep}$ | 0.35 [sec^-1^] | Depolymerization rate of cytoplasmic F-actin |
| $k_{fc1}$ | 8 [sec^-1^] | Disassembly rate of F-actin by cofilin |
| $\alpha$ | 40 [] | Amplification of polymerization of F-actin due to mDia |
| $k_{cn}$ | 0.1 [sec^-1^] | YAP/TAZ nuclear translocation rate independent of F-actin |
| $k_{cy}$ | 20 [sec^-1^] | YAP/TAZ nuclear translocation rate dependent on F-actin |
| $k_{nc}$ | 3 [sec^-1^] | YAP/TAZ cytoplasmic translocation rate independent of LATS_0_ |
| $k_{ly}$ | 6 [sec^-1^] | YAP/TAZ cytoplasmic translocation rate dependent on LATS_0_ |
| $l_{0}$ | 0.5 | Total amount of LATS, LATS_0_ |
| $l_{p}$ | 0.05 | Phosphorylated LATS |
| $\varphi_{0}$*,* $\rho_{0}$*,* $\omega_{0}$*,* $\sigma_{0}$*,* $M_{0}$*,* $L_{0}$*,* $C_{0}$*,* $F_{0}$*,* $Y_{0}$ | 1 | Total amounts of available protein |
| $\lambda$ | 0.1 [] | Maximum inhibitory effect of $Y_{N}$ on DLL4 production |
| $Y_{Y0}$ | 0.47 [cu] | Fraction of $Y_{N}$ from which onwards $Y$ starts inhibiting DLL4 |
| $n$ | 6.5 [] | Scaling the effect of YAP/TAZ inhibition of DLL4 |
| $a$ | -0.6121 [] | Scaling the effect of YAP/TAZ inhibition of LFng |
| $b$ | 1.0578 [] | Scaling the effect of YAP/TAZ inhibition of LFng |
| $n_{1}$ | 0.6 [] | Adapted Hill-function exponent to capture the FAK response in endothelial cells |
| $n_{2}$ | 1.7 [] | Exponent to capture the nonlinear interaction between VEGFR2 and FAK |
| $C_{\varphi}$ | 4.7 [kPa] | $V.R2*E$ value for which activation rate equals half the maximum activation rate |
| $k_{1_{1}}$ | 0.3 [cu^-1^ sec^-1^] | Rate of V.R1 association |
| $k_{{-1}_{1}}$ | 0.001 [sec^-1^] | Rate of dissociation of V.R1 |
| $ɸ_{R_{1}}$ | 0.01 [sec^-1^] | Protein degradation of VR1 |
| $\gamma_{R_{1}}$ | 0.5 [cu sec^-1^] | Protein production of VR1 |
| $k_{up1}$ | 0.0044 [cu^-2^ sec^-1^] | Scales the impact of upregulation of VR1 by H |
| $B_{eff1}$ | 0.62 [] | Scaling factor of effect of ALK1 KO on LFng |
| $B_{eff2}$ | 0.5 [] | Scaling factor of effect of ALK1 KO on Hes/Hey production |

“cu” stands for concentration units.

1.	Passier M, Bentley K, Loerakker S, Ristori T. YAP/TAZ drives Notch and angiogenesis mechanoregulation in silico. npj Syst Biol Appl. 2024 Oct 5;10(1):1–16.

2.	Venkatraman L, Regan ER, Bentley K. Time to Decide? Dynamical Analysis Predicts Partial Tip/Stalk Patterning States Arise during Angiogenesis. PLOS ONE. 2016 Nov 15;11(11):e0166489.

3.	Ristori T, Thuret R, Hooker E, Quicke P, Lanthier K, Ntumba K, et al. Bmp9 regulates Notch signaling and the temporal dynamics of angiogenesis via Lunatic Fringe [Internet]. bioRxiv; 2023 [cited 2024 Jul 16]. p. 2023.09.25.557123. Available from: https://www.biorxiv.org/content/10.1101/2023.09.25.557123v1

4.	Sun M, Spill F, Zaman MH. A computational model of YAP/TAZ mechanosensing. Biophys J. 2016;110(11):2540–50.
